# Gene regulatory co-expression networks decipher potential lncRNA-miRNA-mRNA interactions modulating transcription regulation in neurodegeneration

**DOI:** 10.64898/2026.07.03.736295

**Authors:** Amrit Venkatesan, Prashasti Sinha, Jolly Basak, Ranjit Prasad Bahadur

## Abstract

Neurodegenerative diseases are complex disorders characterised by progressive neuronal loss and widespread transcriptomic dysregulation; however, the coordinated interactions among coding and non-coding RNAs that contribute to disease progression remain incompletely understood. In this study, RNA-seq datasets from disease-relevant neuronal populations and brain regions representing Alzheimer’s disease (AD), Parkinson’s disease (PD) and amyotrophic lateral sclerosis (ALS) were analysed using an integrative network-based framework. Differential expression analysis coupled with weighted gene co-expression network analysis identified modules significantly correlated with disease and prioritised highly connected hub genes. Integration of these hub genes with curated RNA interaction database enabled the construction of candidate lncRNA-miRNA-mRNA regulatory networks. Functional enrichment analysis revealed Gene Ontology biological processes associated with synaptic signalling, mitochondrial function, RNA metabolism and neuroinflammatory responses across neurodegenerative conditions. The inferred regulatory networks suggested both disease-specific and shared post-transcriptional regulatory modules involving key hub genes and non-coding RNAs. Additionally, putative sequence variants were identified within untranslated regions of selected hub genes, suggesting potential alterations in miRNA-mediated regulations. Therefore, this study provides a systems-level view of transcriptomic dysregulation across major neurodegenerative diseases and identifies candidate regulatory interactions and molecular targets for future functional investigation.

## 1. INTRODUCTION

Neurodegenerative disorders are characterised by the progressive loss of selective population of neurons accompanied by cellular inclusions, extracellular protein deposits and morphological abnormalities (Armstrong et al. 2005). The pathological cascade of events underlying neurodegeneration involves a complex interplay of oxidative stress and mitochondrial dysfunction (Mattson 2000). Neurodegenerative diseases can be predominantly categorised based on their principal molecular abnormality, anatomic distribution of neurodegeneration (such as frontotemporal degeneration, extrapyramidal disorders or spinocerebellar degeneration) and primary clinical features including dementia, parkinsonism or motor neuron disease (Dugger and Dickson 2017). The three major neurodegenerative diseases are Alzheimer’s Disease (AD), Parkinson’s Disease (PD) and Amyotrophic Lateral Sclerosis (ALS) (Checkoway et al. 2011). AD, the most common form of dementia, accounts for up to 60% to 80% of overall cases. The underlying pathogenic mechanisms in AD include the amyloid-beta (Aβ) aggregation and tau hyperphosphorylation with tangle formation (Blennow et al. 2006). AD can be predominantly categorised into familial AD (FAD) and sporadic AD (SAD). FAD typically occurs in populations of age below 65 years and accounts for less than 5% of AD cases (Tanzi et al. 1996; Wu et al. 2012). Mutations in three distinct genes, Presenilin 1 (*PSEN1*), Presenilin 2 (*PSEN2*) and Amyloid Precursor Protein (APP) are recognised as causative factors of FAD (Cruts et al. 1998; Hellström-Lindahl et al. 2009; Wu et al. 2012). Sporadic AD accounts for more than 95% of all AD cases and arises from a complex interaction of genetic, environmental and age-associated factors. Among the known genetic risk factors, the APOE [4 allele is the strongest susceptibility factor for late-onset AD; however, it is neither essential nor sufficient to cause disease (William Rebeck et al. 1993; Lippa et al. 1996). PD is characterised by the accumulation of alpha-synuclein-positive Lewy bodies, resulting in progressive loss of dopaminergic neurons primarily in the substantia nigra of the brain stem (Samii et al. 2004; Bloem et al. 2021). Mutations leading to the accumulation of alpha-synuclein are found in genes including *SNCA*, *LRRK2* and *GBA* (Bloem et al. 2021). Jean Martin Charcot originally characterised ALS in 1869 (Rowland 2001). ALS, also known as the “Lou Gehrig’s disease”, is characterised by the progressive loss of the upper motor neurons (UMN) and the lower motor neurons (LMN) (Parkinson et al. 2006; Feldman et al. 2022; Akçimen et al. 2023). ALS typically affects populations above 65 years of age. About 60% of cases of familial ALS (fALS) and 10% of cases of sporadic ALS (sALS) are linked to pathogenic variations of the genes superoxide dismutase 1 (*SOD1*), TAR DNA-binding protein (*TARDBP*) and chromosome 9 open reading frame 72 (*C9orf72*) (Akçimen et al. 2023).

microRNAs (miRNAs) are one of the key players in post-transcriptional gene regulation, which influence various physiological and biochemical processes (Li et al. 2023). They are single-stranded, non-coding RNAs (ncRNAs) with an approximate length of 21 to 24 nucleotides (nt) and are derived from the precursor-miRNAs. miRNAs repress gene transcription by forming RNA-induced silencing complex (RISC) with the Argonaute family of proteins. miRNAs function as key regulators of neuroinflammation in various neurodegenerative disorders. miR-7 and miR-205 regulate *SNCA* and *LRRK2* genes that are involved in etiopathogenesis of PD (Strafella et al. 2018). miR-19b is downregulated approximately five years prior to the onset of motor symptoms in PD (Strafella et al. 2018). An overall downregulation of miR-15b-5p, miR-21-5p and miR-122-3p is reported in cerebrospinal fluid (CSF) of ALS patients (Benigni et al. 2016). miR-107 and miR-29b targeting APP are significantly downregulated in AD (Alshalalfa 2012; Pichler et al. 2017; Goel et al. 2026). Long non-coding RNAs (lncRNAs), on the other hand, are the ncRNAs with length greater than 200 nts involved in transcription regulation. In addition, they function as signals, guides, decoys and scaffolds. Like mRNAs, lncRNAs are majorly transcribed by RNA polymerase II (Wang and Chang 2011). Based on the loci of origin, lncRNAs can be classified as long intergenic non-coding RNA (lincRNA) and lncRNA, where the former is derived from the intergenic regions and the latter is derived from the intronic regions or overlapping exonic regions within the protein-coding genes (Ma et al. 2013). Several lncRNAs contain microRNA response elements (MREs) with partial complementarity to miRNAs, enabling them to act as miRNA sponges, thus preventing their interactions with mRNAs. Such lncRNAs are referred to as competitive endogenous RNAs (ceRNAs). ceRNAs sponging for various miRNAs are crucial in the progression of neurodegenerative disorders. The lncRNA *BACE-AS1* sponges miR-29, targeting *BACE1*, thereby increasing the amyloid production in AD (Goel et al. 2026). *SNHG1* stabilises *SNCA* gene expression by sponging miR-7, resulting in α-synuclein production and accumulation (Thangavelu et al. 2024). Similar ceRNA-mediated regulatory mechanisms have also been reported in other neurodegenerative disorders such as Huntington’s Disease (Ghafouri-Fard et al. 2022). Sponging of miR-129-5p by the lncRNA *NR3C* hinders the suppression of p53 in ALS, which triggers USP10-mediated p53 activation (Pang et al. 2024).

Differential gene expression studies performed using microarrays and RNA-Seq data provide information on relative expression of genes under various conditions. Gene functionalities and gene interactions can be inferred from co-expression patterns using several statistical techniques. Different correlation methods are available for determining the interdependence of two genes, including the Pearson correlation, Spearman correlation, Biweight mid-correlation, Kendall correlation and Distance correlation (de Siqueira Santos et al. 2014). In recent years, machine learning approaches combined with statistical methods have emerged for identifying gene interactions to build gene regulatory networks. Primarily used approaches include Weighted Gene Co-expression Network Analysis (WGCNA), Algorithm for the Reconstruction of Accurate Cellular Networks (ARACNE) and GENIE3 (Margolin et al. 2006; Langfelder and Horvath 2008; Huynh-Thu and Geurts 2018). WGCNA uses an unsupervised approach to cluster genes into modules based on their expression correlation.

Most computational investigations in neurodegeneration analyse differential expression, prediction of lncRNA-miRNA-mRNA network, functional enrichment analysis and variant discovery independently, limiting systems-level interpretation. Predicted lncRNA-miRNA interactions are typically based on sequence complementarity and expression correlation, while the combined effects of expression dysregulation and genetic variation on these networks remain incompletely understood. In particular, the influence of variants within untranslated regions (UTRs) on miRNA binding and gene suppression have only been limitedly integrated with network-based analyses. Present study adopts an integrative computational framework to investigate regulatory mechanisms underlying neurodegenerative diseases. Differential expression analysis is performed to identify significantly dysregulated mRNAs and lncRNAs associated with neurodegeneration. Co-expressed lncRNA-mRNA pairs are incorporated into lncRNA-miRNA-mRNA interaction networks to explore post-transcriptional regulatory mechanisms. Network analysis is further used to prioritise key hub regulators that play central roles in disease-associated regulatory processes. UTR variants are systematically mapped onto the miRNA binding sites of the identified hub genes to evaluate their potential regulatory impact. A schematic representation of the workflow is shown in Fig. 1. This integrated strategy facilitates the identification of altered regulatory network topology, perturbations in functional pathways and mutations that may disrupt miRNA-mediated gene suppression in neurodegeneration.

**Fig. 1.**
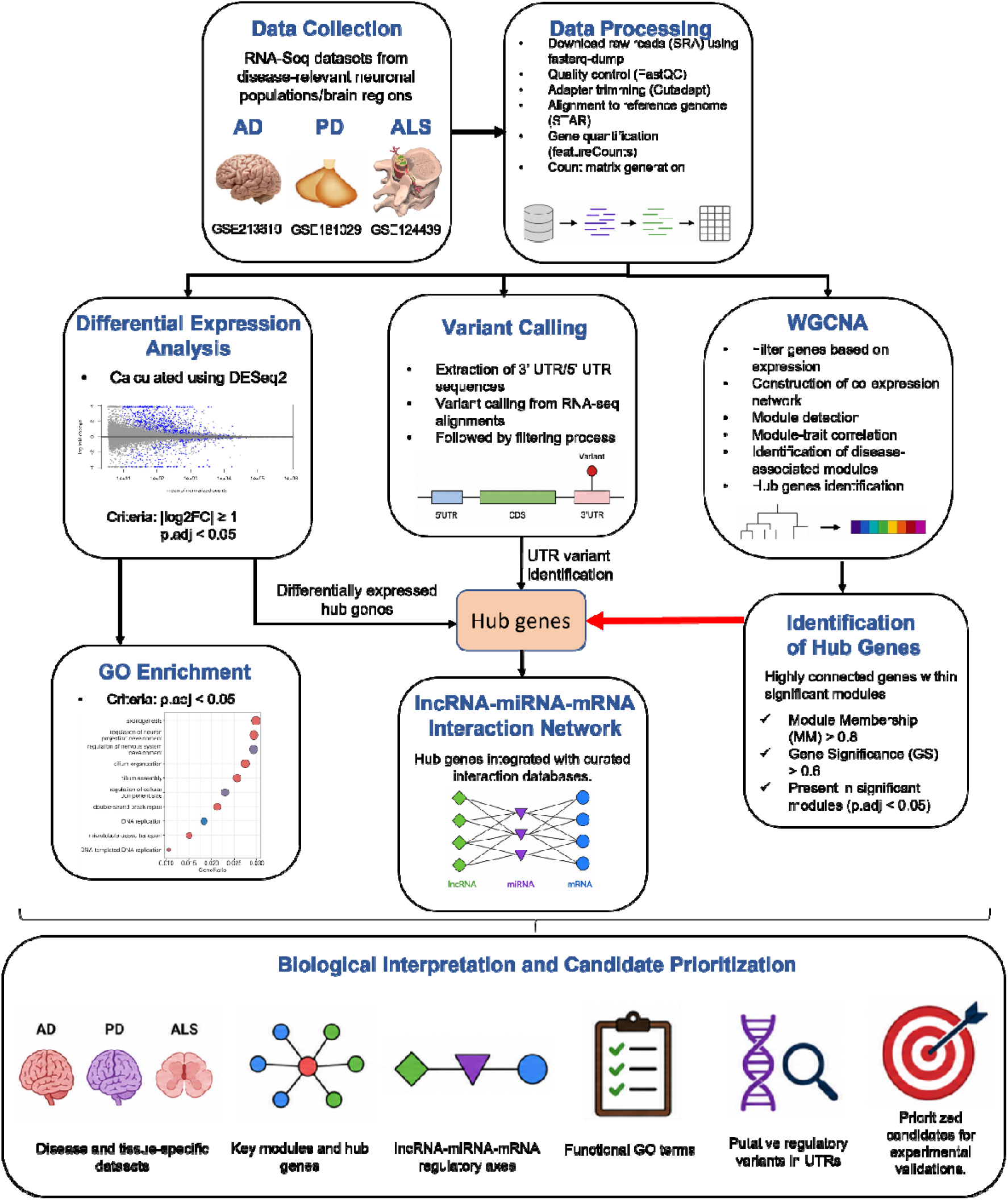
Schematic representation of the study workflow.

## 2. METHODS

### 2.1 RNA-Seq data retrieval and differential gene expression (DGE) analysis

RNA-Seq data for three independent Gene Expression Omnibus (GEO) datasets of AD (GSE213610), PD (GSE181029) and ALS (GSE124439) were analysed in this study (Tam et al. 2019; Brown et al. 2022; Ma et al. 2022; Novosadova et al. 2022; Scopa et al. 2023). The AD dataset contains 32 samples of FAD, SAD and non-AD individuals. The samples were taken from CA3 neurons of the hippocampus and the hippocampal neural precursor cells (hpNPCs). We have eight samples of each subtype and four controls for hpNPCs. For CA3 neurons, we have four samples of each subtype and four controls. The PD dataset contains 36 samples obtained from neural progenitors and terminally differentiated neurons. Each cell type is represented by nine PD and nine control samples. The ALS GEO dataset contains 176 samples categorised as “ALS Spectrum MND”, “Non-neurological Controls” and “Other neurological disorders”. The samples labelled as “Other neurological disorders” were excluded. Finally, 162 samples were retained for downstream analysis. Raw sequence reads for all datasets were obtained from the Sequence Read Archive (SRA). The reads were converted into FASTQ format using the fasterq-dump. Quality assessment was performed using FastQC followed by trimming using cutadapt. The trimmed reads were aligned with human reference genome (GRCh38.p14) using STAR aligner v2.7.10, and gene read counts were generated with Subread v2.0.3 package. DESeq2 v1.34.2 was used to analyse differential gene expression within the R environment. Subsequently, the gene-level count data were analysed using a generalized linear model based on the negative binomial distribution. In order to assess the expression differences between the two groups, disease versus control status was included as the experimental factor in the designed formula. Before fitting the statistical model for each gene, DESeq2 internally estimated size factors for normalization and dispersion parameters. The Wald test and p values were adjusted to evaluate the differential expression by applying Benjamini-Hochberg procedure to control the false discovery rate (FDR). Transcripts with an adjusted p-vales (p.adj) ≤ 0.05 were considered significantly differentially expressed and were classified as differentially expressed lncRNAs (DElncRNAs) or mRNAs (DEmRNAs).

### 2.2 Weighted gene co-expression network analysis

Co-expression analysis of RNA-Seq dataset was performed using WGCNA algorithm v1.72. It calculates a similarity measure *s_ij_*, which is the absolute value of correlation coefficient between the expression profiles of all pairs of genes as shown below (Langfelder and Horvath 2008):

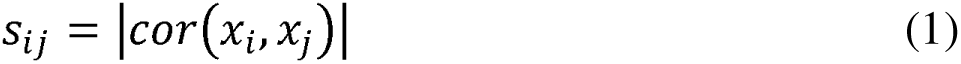

*s_ij_* is used to construct a similarity matrix comprising adjacency values between two genes. The scale-free topology fit indices for different powers can be calculated using pickSoftThreshold function of WGCNA package. The power for which the scale-free topology fit index is greater than 0.8 is considered as the optimal threshold power.

The similarity matrix was used to construct a scale-free network, which defines the gene co-expression and provides highly interconnected hub genes (nodes). The genes were clustered into co-expression modules based on their topological overlap similarity using topology overlap matrix (TOM) in combination with dynamic tree cut algorithm. For each module, a module eigengene (ME) representing the initial principal component analysis (PCA) to characterise the module expression pattern across each sample was computed. Furthermore, the correlation coefficient between the gene and the ME was calculated. Both ME and module membership (MM) were used to assess their association with the characteristic trait of interest. Consequently, the modules having a significant association to the trait are used to construct the scale-free network.

### 2.3 Construction of protein-protein interaction network and lncRNA-miRNA-mRNA network

Protein-protein interaction networks (PPI) were constructed using mRNA genes present in the modules, which are significantly correlated to the traits using the STRING database (Mering 2003). The hub genes of the networks were selected using CytoHubba in Cytoscape v3.10 (Chin et al. 2014). CytoHubba ranks nodes based on multiple topological algorithms, and hub genes were selected according to the maximum clique centrality (MCC) metric. lncRNAs co-expressed with the hub genes were extracted from each significant module. The lncRNA-miRNA and mRNA-miRNA interactions were obtained from NPInter v5.0 and RNAInter v4.0 databases, which comprise of both experimentally validated interactions and interactions predicted using tools such as TargetScan, miRDB and miRTarBase (Wang 2008; Friedman et al. 2009; Hsu et al. 2011; Kang et al. 2022; Zheng et al. 2023). miRNAs commonly sharing interactions between the co-expressed lncRNAs and mRNAs were identified to construct the lncRNA-miRNA-mRNA networks.

### 2.4 Gene ontology enrichment analysis

Gene ontology (GO) enrichment analysis was performed using clusterProfiler package v4.6.2 in R for both the differentially expressed genes and the genes present in the modules significantly associated with the diseases. The biological processes defined by GO terms having enrichment score with p.adj ≤ 0.05 calculated using Benjamini-Hochberg procedure were considered significant. The top enriched GO terms for each disease were subsequently identified. These enriched terms were further analysed to understand the key biological processes. Visualisation of the enriched GO biological processes was performed to highlight the most significantly overrepresented biological processes across the different neurodegenerative diseases.

### 2.5 Variant calling analysis

Variant calling was performed using bcftools v1.13. The BAM files generated using the raw RNA-Seq reads of the AD, PD and ALS datasets were used as an input to obtain genetic variants in each sample. Variants with a quality score exceeding 30 were selected for downstream analysis and subsequently annotated using ANNOVAR tool (2020Jun07 release) (Wang et al. 2010). Mutations present in different genomic regions of hub genes were systematically catalogued across all three diseases. Target genes, their transcript loci and their corresponding miRNA sequences were obtained from miRWalk 3.0 (Sticht et al. 2018). Transcript-level coordinates were converted to genomic coordinates using pyEnsembl for Ensembl release (v113). Mutations specific to miRNA target sites in 3’ UTRs and 5’ UTRs were identified. Moreover, putative novel mutations were identified by cross-referencing the detected mutations against multiple variant databases including dbSNP, gnomAD, ClinVar and 1000 Genomes (Sherry 2001; Landrum et al. 2014; Devuyst 2015; Karczewski et al. 2020). Candidate putatively novel UTR variants were subsequently evaluated using supplementary quality metrics, including read depth, mapping quality, allele fraction, and strand support, and only those exhibiting consistent evidence across these metrics were considered high-confidence candidates.

## 3. RESULTS

### 3.1 Differentially expressed mRNA and lncRNA in different tissues of AD, PD and ALS

The obtained raw read count matrices contain 47363 genes of AD, 50218 genes of PD and 27549 genes of ALS which includes both the lncRNA and mRNA genes. Genes having low read counts below 30 across all samples are excluded from further analysis. Following filtering, 16383 genes of AD (GSE213610), 14469 genes of PD (GSE181029) and 15941 genes of ALS (GSE124439) are retained. PCA was performed on variance-stabilized expression values to assess sample clustering and identify potential outliers prior to downstream analyses. Filtered genes are subjected to differential gene expression analysis using DeSeq2 package in R, which uses median-of-ratios normalisation. Genes encoding mRNAs and lncRNAs having |log2 fold change (FC)| > 1.0 and with a false discovery rate adjusted p.value ≤ 0.05 are considered as differentially expressed. Total number of DEmRNAs and DElncRNAs in different tissues of AD, PD and ALS are listed in Table 1.

**Table 1:** The number of differentially expressed mRNAs and lncRNAs in AD, PD and ALS.

| Disease | Tissue type |  | DEmRNAs | DElncRNAs |
| --- | --- | --- | --- | --- |
| AD | CA3 neurons | FAD | 1058 (70% (↑) & 30 % (↓)) | 42 (86% (↑) & 14% (↓)) |
|  |  | SAD | 68 (32% (↑) & 68% (↓)) | 2 (00% (↑) & 100% (↓)) |
|  | hpNPCs | FAD | 1872 (71% (↑) & 29% (↓)) | 238 (88% (↑) & 12% (↓)) |
|  |  | SAD | 149 (81% (↑) & 19% (↓)) | 19 (74% (↑) & 26% (↓)) |
| PD | Neural progenitors |  | 3080 (62% (↑) & 38% (↓)) | 357 (52% (↑) & 48% (↓)) |
|  | Terminally differentiated neurons |  | 3021 (54% (↑) & 46% (↓)) | 300 (38% (↑) & 62% (↓)) |
| ALS | Frontal cortex |  | 1943 (38% (↑) & 62% (↓)) | 187 (10% (↑) & 90% (↓)) |
|  | Motor cortex |  | 1606 (41% (↑) & 59% (↓)) | 110 (26% (↑) & 74% (↓)) |

(↑) and (↓) represent percentage of upregulated and downregulated differentially expressed genes, respectively.

The distribution of differentially expressed genes in different tissues of AD, PD and ALS is shown in Fig. 2. In SAD, both CA3 neurons and hpNPCs show fewer differentially expressed genes. A higher percentage of differentially expressed genes are downregulated in both the frontal and motor cortices of ALS (Table 1). The AD cohort contains both FAD and SAD patient samples from two different tissue types, the hippocampal CA3 neurons and hpNPCs. A total of 1058 mRNAs and 42 lncRNAs are differentially expressed in CA3 neurons in FAD, whereas SAD samples exhibit differential expression of 68 mRNAs and 2 lncRNAs. For hpNPCs, 1872 mRNAs and 238 lncRNAs are differentially expressed in FAD, while 149 mRNAs and 19 lncRNAs are differentially expressed in SAD (Supplementary Tables S1 and S2). The PD dataset contains patient samples obtained from neural progenitors and terminally differentiated neurons. Neural progenitors in PD exhibit 3080 differentially expressed mRNAs and 357 differentially expressed lncRNAs. On the other hand, 3021 mRNAs and 300 lncRNAs are differentially expressed in terminally differentiated neurons (Supplementary Tables S1 and S2). The dataset of ALS contains patient samples from the frontal cortex and motor cortex of the brain. Differential expression analysis identified 1943 mRNAs and 187 lncRNAs in the frontal cortex, and 1606 mRNAs and 110 lncRNAs in the motor cortex (Supplementary Tables S1 and S2).

**Fig. 2.**
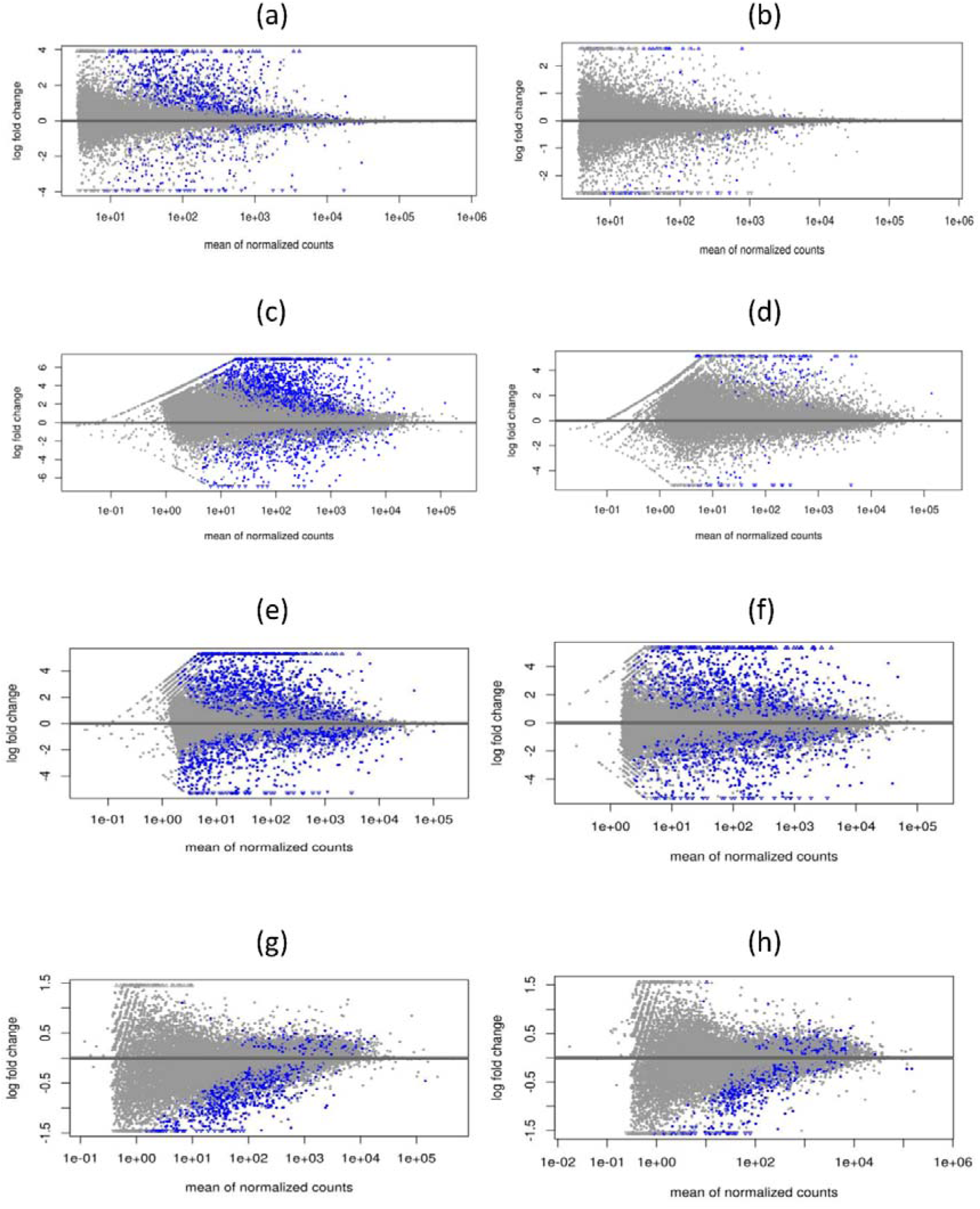
MA plots showing distribution of gene expression in (a) AD (CA3 neurons), (b) AD (hpNPCs), (c) PD (neural progenitors), (d) PD (terminally differentiated neurons), (e) ALS (frontal cortex), (f) ALS (motor cortex). The blue dots represent the differentially expressed genes.

### 3.2 Identification of co-expression modules of mRNA and lncRNA

WGCNA analysis for different tissue types are performed independently for each dataset. Outlier samples are identified and removed using hierarchical clustering. Biweight mid-correlation matrices are constructed using the retained genes and subsequently transformed into adjacency matrices for each dataset. TOM-based dissimilarity measure in conjunction with dynamic cut tree algorithm reveals the genes clustered into different modules. The cluster dendrogram for AD, PD and ALS is displayed in Fig. 3. The construction of networks and module detection are performed using blockwiseModules function in WGCNA with maxBlockSize set to the number of genes retained after outlier removal. Module-trait relationships are visualised using a heatmap (Supplementary Fig. S1), and inter-module correlations are assessed using a module eigengene dendrogram heatmap (Supplementary Fig. S2).

**Fig. 3.**
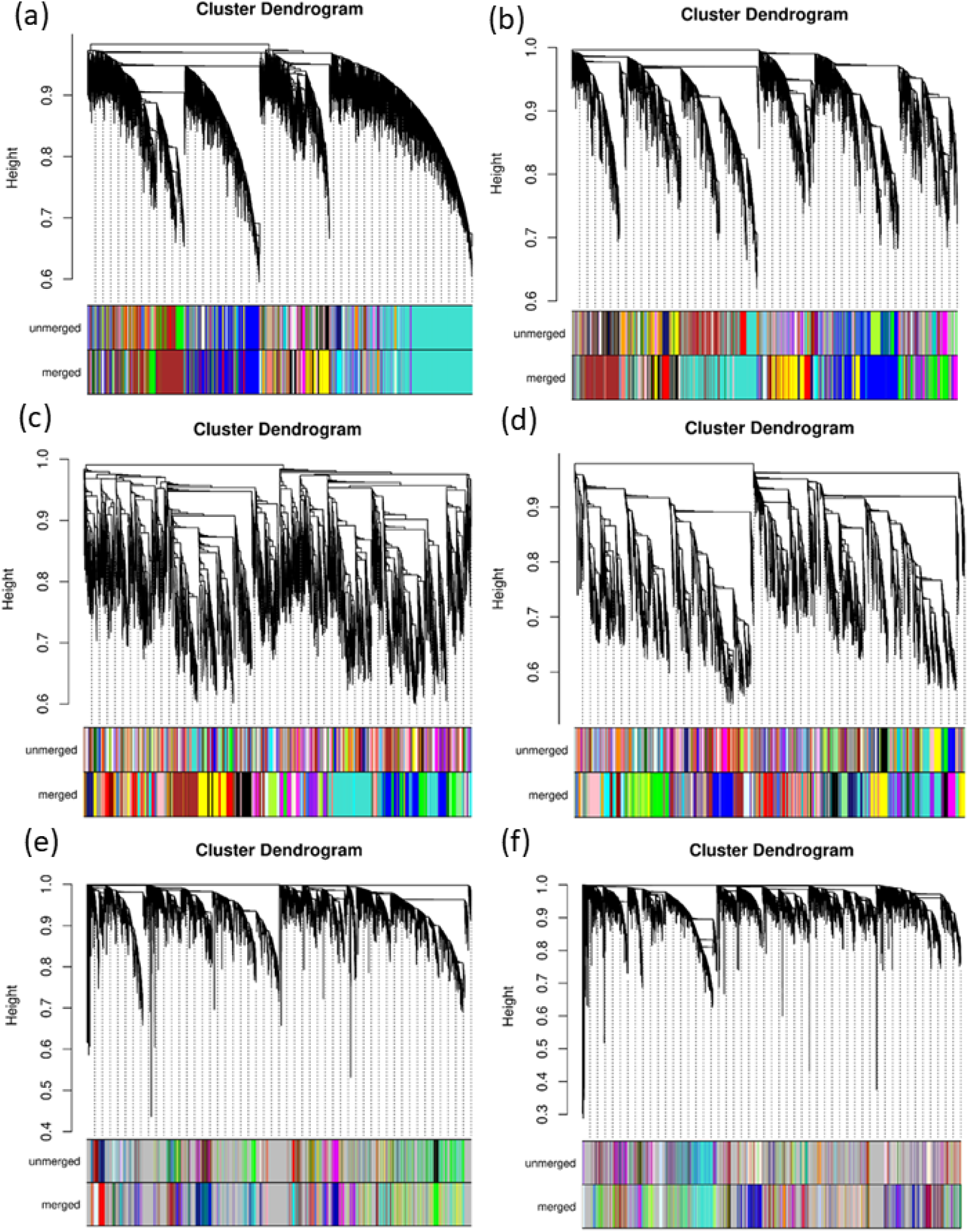
Dendrograms for various modules representing gene clusters in (a) AD (CA3 neurons), (b) AD (hpNPCs), (c) PD (neural progenitors), (d) PD (terminally differentiated neurons), (e) ALS (frontal cortex), (f) ALS (motor cortex).

The adjacency matrix for AD is obtained using soft-threshold power *β* = 8 (*R*² = 0.98) for CA3 neurons and *β* = 14 (*R*² = 0.97) for hpNPCs, ensuring approximate scale-free topology. The genes are clustered into 49 distinct modules for CA3 neurons and 24 distinct modules for hpNPCs (Supplementary Fig. S1). The modules coloured in blue, red, green and light-yellow (p.val ≤ 0.05) are significantly associated to FAD (Supplementary Fig. S1(a)), while the modules in sky-blue, royal-blue, floral-white and white show significant association with SAD in CA3 neurons (Supplementary Fig. S1(b)). Similarly, the modules in light-cyan, red, midnight-blue, dark-green, pink, black and light-yellow are significantly associated to FAD (Supplementary Fig. S1(c)). Although the grey module showed significant correlation, it was excluded from downstream biological interpretation because it represents genes that do not belong to a well-defined co-expression module (Supplementary Fig. S1(d)). In the PD dataset, soft-threshold power *β* = 12 (*R*² = 0.97) for neural progenitors and *β* = 8 (*R*² = 0.94) for terminally differentiated neurons are selected to achieve scale-free topology. WGCNA identifies 35 co-expression modules in neural progenitors and 28 modules in terminally differentiated neurons (Supplementary Fig. S1). In neural progenitors, light-cyan, green-yellow, magenta, brown, black and violet modules exhibit significant correlation with PD (Supplementary Fig. S1(e)). On the other hand, midnight-blue, dark-turquoise, salmon, light-green, grey, dark-grey and black modules are significantly correlated with PD in terminally differentiated neurons (Supplementary Fig. S1(f)).

In ALS dataset, adjacency matrices are constructed using soft-threshold power *β* = 18 (*R*² = 0.90) for frontal cortex samples and *β* = 14 (*R*² = 0.94) for motor cortex samples. WGCNA identifies 24 modules in frontal cortex tissues and 48 modules in motor cortex tissues (Supplementary Fig. S1). Among these, the yellow module shows a significant correlation with ALS in the frontal cortex (Supplementary Fig. S1(g)), while the midnight-blue module is significantly associated with ALS in the motor cortex (Supplementary Fig. S1(h)).

### 3.3 Interaction network among lncRNA-miRNA-mRNA

Following the identification of significant modules in different tissue types of AD, PD and ALS, PPI networks are constructed using the mRNA genes present in the significant modules. Hub genes are selected for different tissue types of AD, PD and ALS using MCC score. They have significant correlation to the trait (p.val ≤ 0.05) and have significant module membership (p.val ≤ 0.05) (Supplementary Table S3). lncRNAs co-expressed with hub genes in the significant modules are also identified. To construct the lncRNA-miRNA-mRNA interaction network, miRNAs interacting with the identified lncRNA-mRNA pairs are retrieved from the NPInter and RNA Inter databases (Fig. 4).

**Fig. 4.**
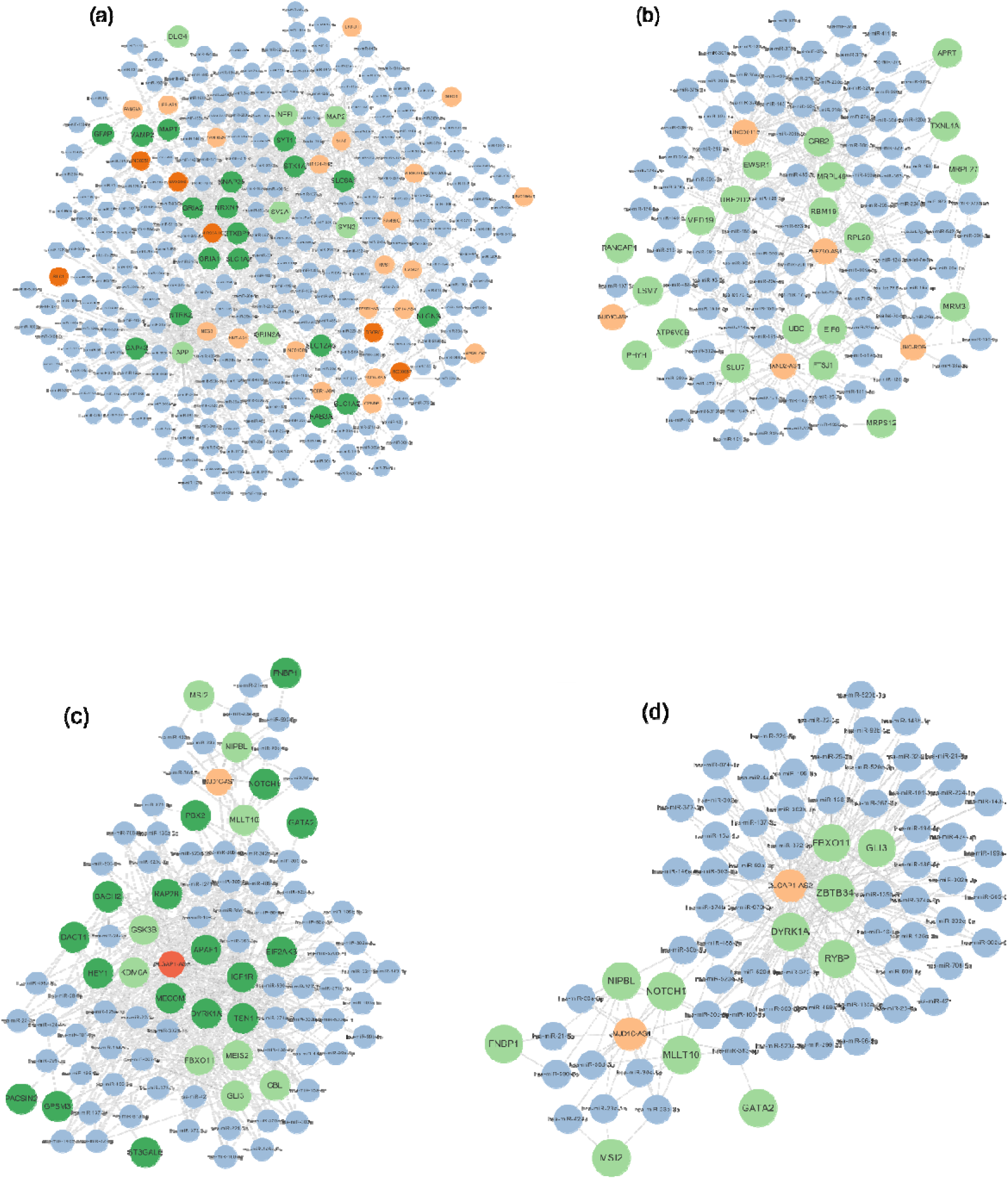
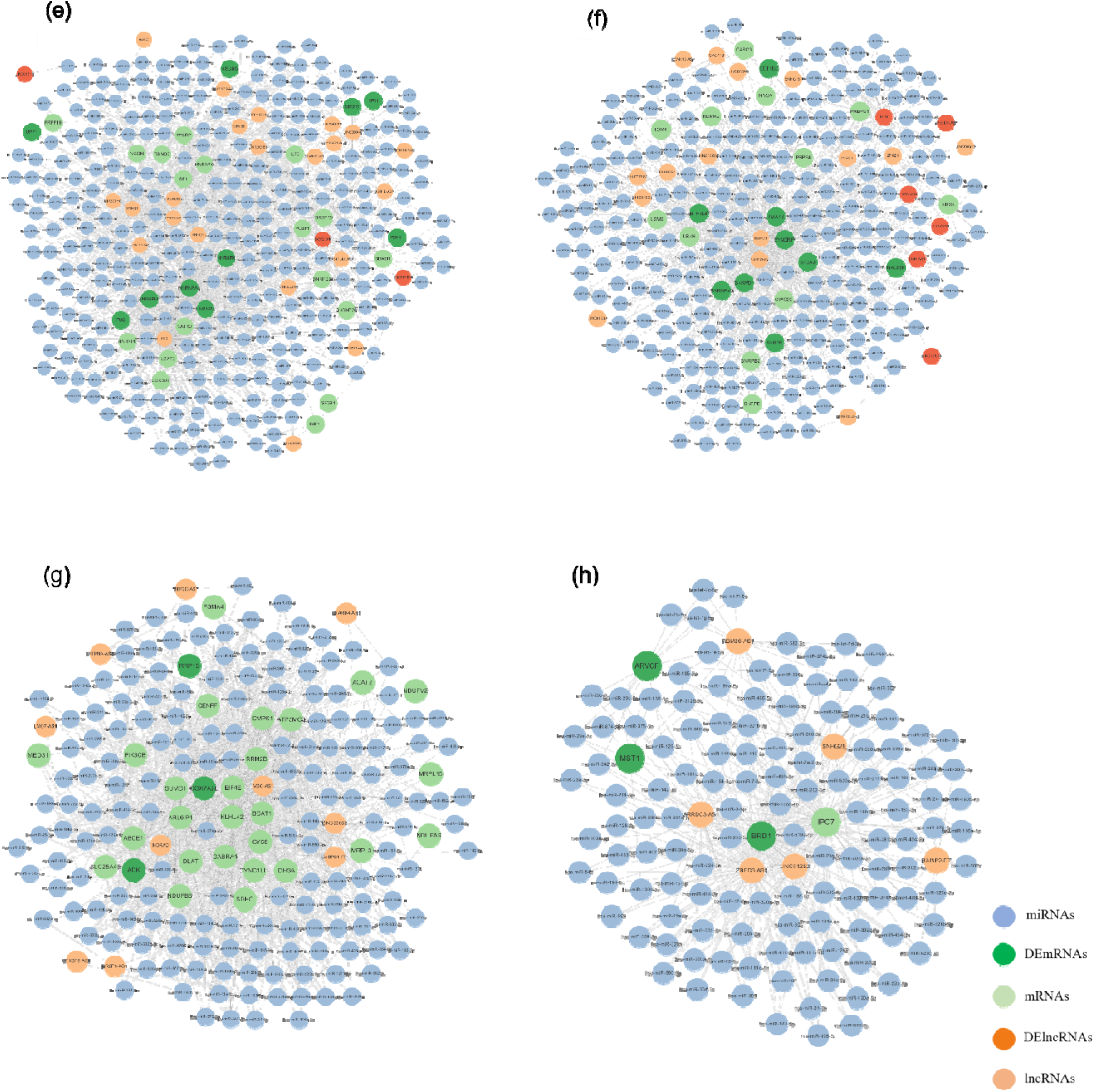
lncRNA-miRNA-mRNA networks in (a) FAD (CA3 neurons), (b) SAD (CA3 neurons), (c) FAD (hpNPCs), (d) SAD (hpNPCs), (e) PD (neural progenitors), (f) PD (terminally differentiated neurons), (g) ALS (frontal cortex), (h) ALS (motor cortex).

Overall number of interactions among lncRNA-miRNA-mRNA in AD, PD and ALS are shown in Table 2. In CA3 neurons of FAD patients, 900 lncRNA-miRNA-mRNA interactions are identified involving 27 lncRNAs, 242 miRNAs and 25 mRNAs. In SAD patients, 230 lncRNA-miRNA-mRNA interactions are identified involving 5 lncRNAs, 95 miRNAs and 20 mRNAs. In hpNPCs of FAD patients, 364 lncRNA-miRNA-mRNA interactions are identified involving 2 lncRNAs, 69 miRNAs and 26 mRNAs. In SAD patients, 172 lncRNA-miRNA-mRNA interactions are identified involving 2 lncRNAs, 66 miRNAs and 11 mRNAs. Neural progenitors in PD patients exhibit 1099 lncRNA-miRNA-mRNA interactions involving 25 lncRNAs, 315 miRNAs and 28 mRNAs. In terminally differentiated neurons, 651 lncRNA-miRNA-mRNA interactions are identified involving 21 lncRNAs, 255 miRNAs and 21 mRNAs. In frontal cortex tissues of ALS, 1191 lncRNA-miRNA-mRNA interactions are identified involving 10 lncRNAs, 166 miRNAs and 29 mRNAs. On the other hand, 262 lncRNA-miRNA-mRNA interactions are identified involving 6 lncRNAs, 123 miRNAs and 4 mRNAs in motor cortex tissues. A detailed list of mRNA, miRNA and lncRNA involved in interaction networks in AD, PD and ALS is provided in Supplementary Table S4. The identified networks highlighted several candidate disease-associated regulatory modules. AD networks contained lncRNAs linked to amyloidogenic pathways and neuronal maintenance, whereas PD networks were enriched for regulators associated with dopaminergic neuronal function and synaptic signalling. ALS-specific networks predominantly involved regulators associated with RNA metabolism, cellular stress responses and neuronal survival pathways. These observations suggest both disease-specific and partially overlapping post-transcriptional regulatory programs across neurodegeneration.

**Table 2:** Number of lncRNA-miRNA-mRNA interactions for the hub genes identified in AD, PD and ALS.

| Disease | Tissue type |  | lncRNA-miRNA-mRNA interactions<sup>a</sup> |
| --- | --- | --- | --- |
| AD | CA3 neurons | FAD | 900 |
|  |  | SAD | 230 |
|  | hpNPCs | FAD | 364 |
|  |  | SAD | 172 |
| PD | Neural progenitors |  | 1099 |
|  | Terminally differentiated neurons |  | 651 |
| ALS | Frontal cortex |  | 1191 |
|  | Motor cortex |  | 2621 |

<sup>a</sup>Number of interactions are identified by calculating the number of edges in the network

### 3.4 Functional enrichment of differentially expressed genes and co-expressed genes in significant modules of AD, PD and ALS

GO enrichment analysis is performed for the differentially expressed genes and co-expressed genes in significant modules of AD, PD and ALS. Table 3 shows the number of enriched GO biological processes for differentially expressed genes in AD, PD and ALS. In FAD, 700 GO terms are enriched in CA3 neurons, whereas 1027 GO terms are enriched in hpNPCs. In SAD, 55 GO terms are enriched in CA3 neurons, whereas only 3 pathways are enriched in hpNPCs. In PD, 932 GO terms are enriched in neural progenitors, whereas 987 GO terms are enriched in terminally differentiated neurons. In ALS, 8 GO terms in frontal cortex tissues and 8 GO terms in motor cortex tissues are enriched. A higher number of enriched terms in FAD and PD indicate that the differentially expressed genes have a much broader range of biological functions leading to more widespread molecular dysregulation. A detailed list of enriched GO biological processes for the differentially expressed genes and co-expressed genes in significant modules of AD, PD and ALS is provided in Supplementary Table S5 and Table S6, respectively. The top enriched GO biological processes in different diseases are shown in Fig. 5, which indicates the ratio between the number of differentially expressed genes to the total number of genes involved in a particular GO process. Terms associated with axonogenesis, regulation of neuron differentiation, and extracellular matrix organization are consistently enriched in both FAD and PD. This indicates common disruptions in neuronal development, synaptic connectivity, synaptic stability and neuronal repair mechanisms. In contrast, SAD and ALS do not share any common enriched pathways, either with each other or with FAD and PD. Furthermore, no shared enriched pathways are observed between the frontal cortex and motor cortex datasets in ALS or between the CA3 neurons and hpNPCs in SAD, thus indicating a high degree of molecular heterogeneity.

**Fig. 5.**
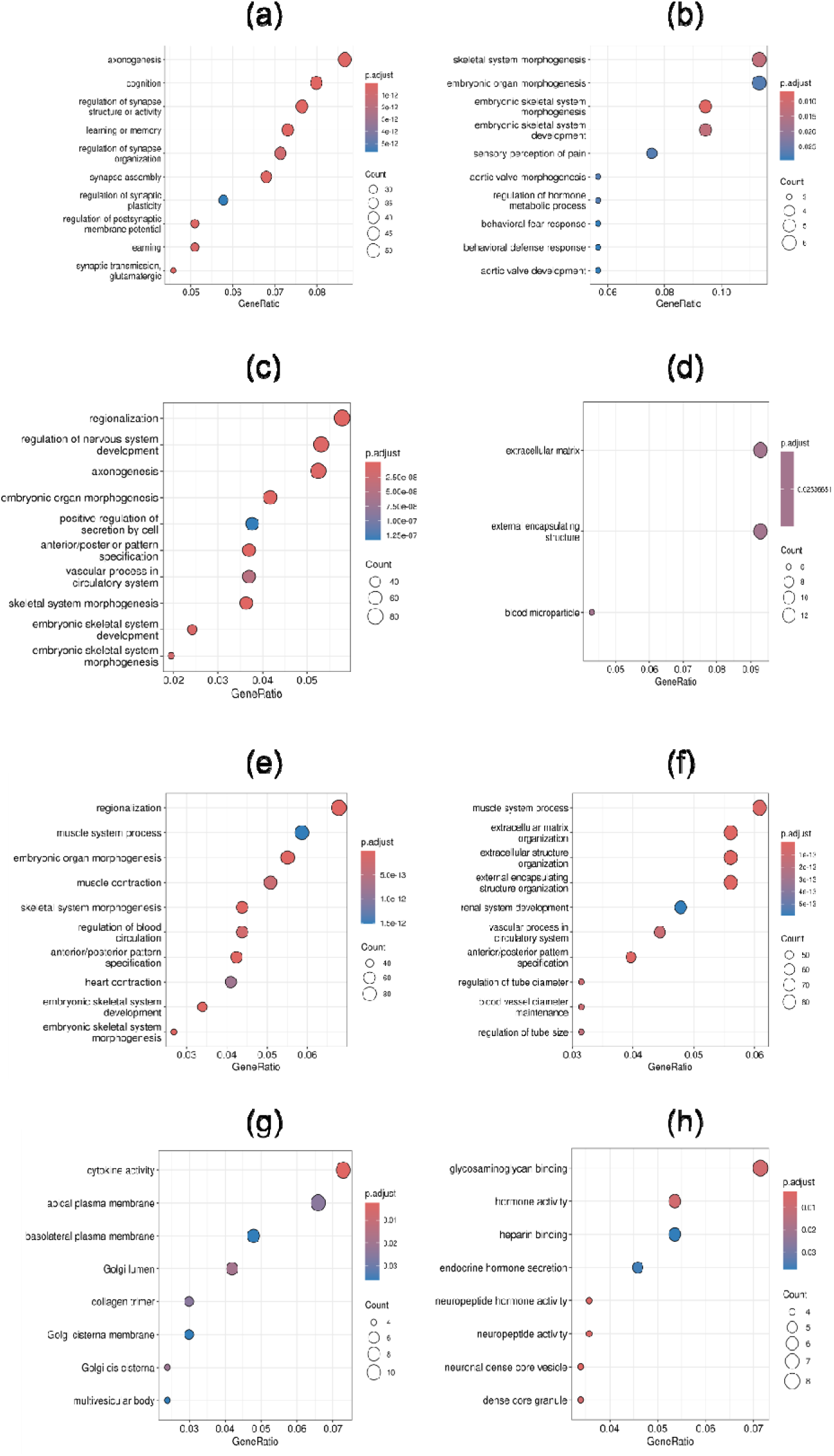
Top ten enriched pathways in (a) FAD (CA3 neurons), (b) SAD (CA3 neurons), (c) FAD (hpNPCS), (d) SAD (hpNPCs), (e) PD (neural progenitors), (f) PD (terminally differentiated neurons), (g) ALS (frontal cortex), (h) ALS (motor cortex).

**Table 3:** Number of significantly enriched pathways in AD, PD and ALS.

| Disease | Tissue type |  | Number of enriched pathways |
| --- | --- | --- | --- |
| AD | CA3 neurons | FAD | 700 |
|  |  | SAD | 55 |
|  | hpNPCs | FAD | 1027 |
|  |  | SAD | 3 |
| PD | Neural progenitors |  | 932 |
|  | Terminally differentiated neurons |  | 987 |
| ALS | Frontal cortex |  | 8 |
|  | Motor cortex |  | 8 |

### 3.5 Variants in different genomic regions of hub-genes in AD, PD and ALS

Variant calling analysis is carried out across all the samples in AD, PD and ALS datasets. To minimise the sequencing errors, only the variants with a high-quality score greater than 30 are considered. The distribution of variants across different genomic regions of hub genes in the various tissue types and disease conditions is summarised in Table 4. The list of complete set of mutations as well as putative novel mutations present in miRNA-interacting regions of 3’ UTRs and 5’ UTRs of the hub genes is tabulated in Supplementary Table S7. The reported putatively novel UTR variants exhibited strong supporting evidence, including high variant quality scores, adequate read depth (DP ≥ 30), sufficient alternate allele support, and acceptable mapping quality (MQ ≥ 20), supporting their classification as high-confidence candidate variants. No novel variants in 3’ UTRs and 5’ UTRs of hub genes are identified in CA3 neurons of FAD and SAD patients, hpNPCs of SAD patients, neural progenitors of PD patients and motor cortex of ALS patients.

**Table 4:** Number of mutations in different genomic regions of hub genes in AD, PD and ALS.

| Disease | AD |  |  |  | PD |  | ALS |  |
| --- | --- | --- | --- | --- | --- | --- | --- | --- |
| Tissue type | CA3 neurons |  | hpNPCs |  | Neural progenitors | Terminally differentiated neurons | Frontal cortex | Motor cortex |
|  | FAD | SAD | FAD | SAD |  |  |  |  |
| intronic | 202 | 128 | 452 | 799 | 42 | 74 | 7258 | 7441 |
| Splicing | 21 | 40 | 29 | 32 | 22 | 23 | 33 | 30 |
| UTR3 | 57 | 32 | 91 | 102 | 43 | 43 | 543 | 530 |
| UTR5 | 9 | 4 | 14 | 16 | 6 | 2 | 24 | 23 |
| Exonic (non-synonymous) | 3 | 13 | 11 | 22 | 10 | 8 | 28 | 28 |
| Exonic (synonymous) | 20 | 16 | 13 | 18 | 22 | 12 | 33 | 30 |

## 4. DISCUSSION

This study integrates differential expression analysis, WGCNA and lncRNA-miRNA-mRNA interaction mapping to investigate transcriptomic dysregulation across AD, PD and ALS. Although these disorders exhibit distinct clinical and pathological manifestations, our analyses reveal convergence on biological processes related to synaptic signalling, RNA metabolism, mitochondrial function and cellular stress responses. The identified co-expression modules and regulatory interaction networks provide a systems-level framework for understanding how coding and non-coding RNAs may collectively contribute to neurodegenerative disease progression. The inferred regulatory relationships should be interpreted as candidate interactions that prioritise genes and non-coding RNAs for future functional investigation rather than definitive mechanistic pathways.

### 4.1 Transcriptomic regulation in Alzheimer’s disease

We observe significant upregulation of *SNAP25* in CA3 neurons of FAD from hiPSC-derived CA3 neurons and hpNPCs of early-onset AD patients. SNAP25 is a core component of the SNARE complex that interacts with *STXBP1* and is essential for vesicle priming and synaptic neurotransmitter release (Hussain 2014). Elevated *SNAP25* levels is detected in CSF during preclinical AD stages, suggesting that its upregulation may represent a compensatory synaptic response preceding neuronal loss (Zhang et al. 2025). Notably, elevation of *SNAP25* is absent in hpNPCs, which reflect their immature neuronal phenotype and incomplete development of synaptic machinery. Further, upregulation of the lncRNA *PANTR1* in differentiating neuronal cells is identified. *PANTR1* is implicated in neuronal differentiation and apoptosis suppression, consistent with its reported interaction with HuR during neuronal stem cell maturation (Carelli et al. 2019). Similarly, *MIR99AHG*, a lncRNA associated with cell differentiation, is significantly upregulated in both CA3 neurons and hpNPCs of FAD. Dysregulation of *MIR99AHG* is also reported in the hippocampus of individuals with Down syndrome, indicating its role in aberrant neurodevelopmental and neurodegenerative pathways (Zhao et al. 2024). *MIR99AHG* and *FAM182A* are known to sponge miR-27b-3p, miR-181a-5p and miR-23a-3p, which suppress *SNAP25* expression. These observations suggest a potential mechanism through which dysregulated lncRNA expression may influence miRNA-mediated regulation of *SNAP25*; however, experimental validation will be required to establish causality.

### 4.2 Transcriptomic regulation in Parkinson’s disease

Neural progenitors in PD show upregulation of genes such as *NGFR*, *FUS* and *PLA2G4A*, suggesting increased neural plasticity, levodopa induced dyskinesia, lipid dysregulation and failure of the clearance of protein aggregate. These findings are in accordance with the previous studies (Wang et al. 2023). Expression of *COMT* is significantly reduced in PD, implying decreased dopamine catabolism and potentially elevated dopamine levels. Our observation correlates with the previous genotype–phenotype studies in PD (Qian et al. 2017). Upregulation of *FUS* is also linked to its mislocalization from nucleus to cytoplasm in neural progenitors. Furthermore, the identification of the lncRNA SNHG1, which potentially regulates KHSRP through the sequestration of multiple miRNAs, suggests perturbations in RNA-processing and post-transcriptional regulatory pathways in terminally differentiated neurons.

### 4.3 Transcriptomic regulation in Amyotrophic Lateral Sclerosis

In spinal cord motor neurons, *MST1* exhibits post-translational activation via ROS-mediated dissociation from thioredoxin-1, driving neurodegeneration independent of transcriptional changes (Lee et al. 2013). In contrast, the motor cortex data of ALS reveals coordinated transcriptional downregulation of *MST1* alongside *GBF1*, *COX7A2L* and *RRP15*, potentially reflecting region-specific alterations in proteostasis-related pathways in upper motor neurons preceding axonal degeneration. Furthermore, the lncRNA *NORAD*, previously implicated in AD and PD, is significantly co-expressed with *RRP15* and *COX7A2L*. Both of these coding genes are associated with oxidative stress response and mitochondrial function, suggesting that NORAD-centred regulatory interactions may contribute to the maintenance of neuronal homeostasis under conditions of cellular stress. The ALS-associated regulatory networks identified in this study highlight the potential involvement of non-coding RNAs in coordinating pathways related to RNA metabolism, mitochondrial function and stress adaptation. The obtained results support growing evidence that dysregulation of RNA-mediated regulatory mechanisms represents a recurring feature of neurodegenerative pathology. Additionally, putative variants were identified within miRNA-interacting untranslated regions of several hub genes, suggesting possible alterations in post-transcriptional regulation. However, the functional consequences of these variants remain to be experimentally validated (Tan et al. 2015; Vainberg Slutskin et al. 2018).

### 4.4 Shared transcriptomic features across neurodegenerative diseases

Despite differences in tissue origin, disease pathology and experimental context, several common biological themes emerged across AD, PD and ALS. Functional enrichment analyses consistently highlighted processes associated with neuronal development, axonogenesis, synaptic organization, intracellular transport and RNA regulatory mechanisms. These results are consistent with growing evidence that neurodegenerative diseases share fundamental disruptions in neuronal maintenance pathways, even when the initiating molecular events differ (Dugger and Dickson 2017). The identification of overlapping functional signatures suggests that perturbations in RNA-mediated regulatory networks may represent a common feature of neurodegeneration. At the same time, the observed similarities should be interpreted cautiously because the analysed datasets originate from distinct tissues and cellular systems. As a result, the present observations are best viewed as candidate shared mechanisms that require validation in harmonised cohorts and experimental models.

### 4.5 Study limitations

Several limitations should be considered when interpreting the findings of this study. First, the analysed datasets were derived from different tissues, cell types and experimental systems, which may introduce biological and technical heterogeneity that cannot be completely disentangled from disease-specific effects. Second, the identified lncRNA-miRNA-mRNA interactions were inferred from curated interaction databases and should therefore be considered candidate regulatory relations. Third, the co-expression modules identified by WGCNA represent statistical associations rather than direct regulatory interactions. Finally, the variant analysis was performed using RNA-Seq data and consequently may be influenced by transcript abundance, RNA editing events and mapping biases. Therefore, the identified variants should be regarded as putative regulatory candidates requiring independent validation using orthogonal genomic approaches.

## 5. CONCLUSION

We present an integrative systems-level analysis of transcriptomic dysregulation across AD, PD and ALS through the combined application of differential expression analysis, WGCNA and lncRNA-miRNA-mRNA interaction mapping. This approach enabled the identification of disease- and tissue-associated co-expression modules, highly connected hub genes and candidate non-coding RNA regulatory interactions that may contribute to neurodegenerative processes. Functional enrichment analyses consistently highlighted biological processes associated with synaptic organisation, neuronal development, RNA metabolism and cellular stress responses, suggesting that distinct neurodegenerative disorders may share common transcriptomic signatures despite their diverse clinical and pathological manifestations. The inferred lncRNA-miRNA-mRNA networks further emphasise the potential importance of RNA-mediated regulatory mechanisms in shaping disease-associated gene expression programs. The identification of putative variants within miRNA-interacting untranslated regions of selected hub genes suggests possible alterations in post-transcriptional regulation, although the functional consequences of these variants require independent validation. The results support a network-centric view of neurodegeneration in which coordinated perturbations of coding and non-coding RNA regulatory circuits contribute to transcriptomic dysfunction. The framework presented here provides a resource for prioritising candidate genes and regulatory RNAs for future experimental studies and may facilitate the discovery of biomarkers and therapeutic targets relevant to neurodegenerative disease biology.

## Statements & Declarations

## Acknowledgment

AV and PS thank IIT Kharagpur for their fellowship. JB thank Visva-Bharati and RPB thank IIT Kharagpur for infrastructure support. Both JB and RPB thank DBT, India for National Network Project.

## Author Contributions

RPB and AV conceptualized the study, AV, RPB and JB designed the study, AV curated the dataset and performed the work. AV, PS, JB and RPB performed data analysis and wrote the manuscript. All authors read and approved the final manuscript.

## Supplementary information

Available as Supplementary Figs. S1-S2 and Supplementary Tables S1-S7.

## Data availability

The source codes used during the current study can be found at https://github.com/amrit-debug/CODEG/tree/main.

## Ethics approval

Not applicable

