## Supplementary Figures for "Gene regulatory co-expression networks decipher potential lncRNA-miRNA-mRNA interactions modulating transcription regulation in neurodegeneration"

(a)

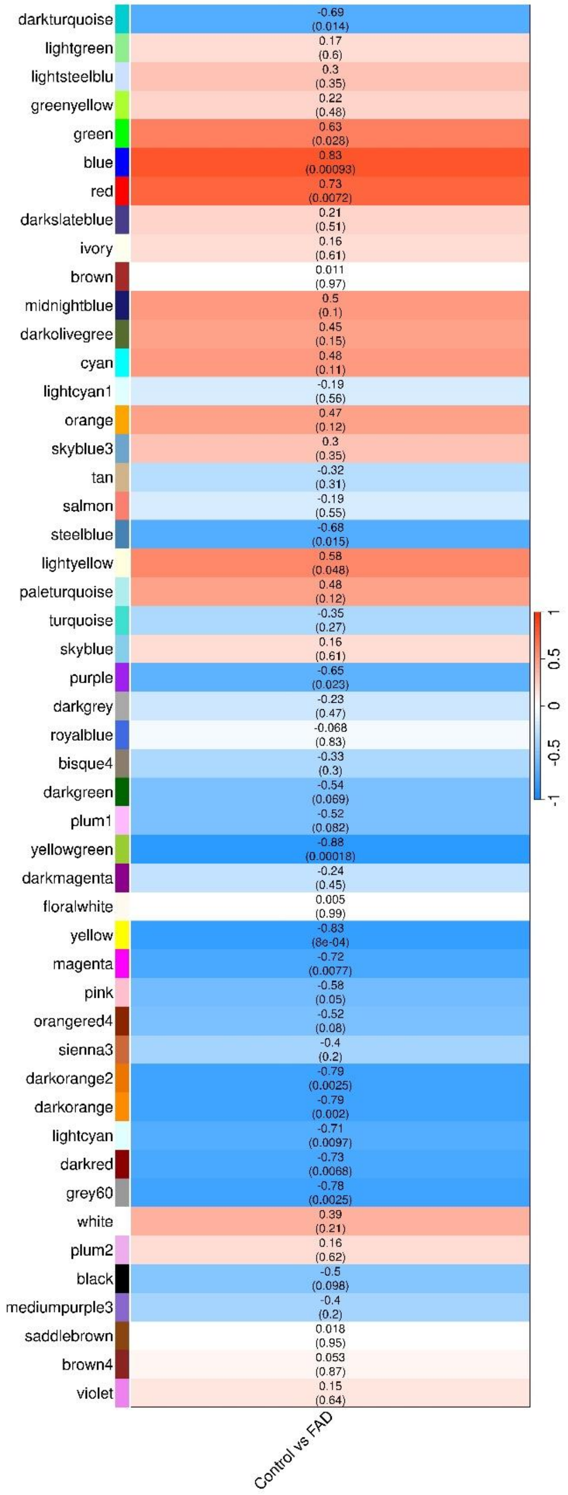
darkturquoise
-0.69
(0.014)
lightgreen
0.17
(0.6)
lightsteelblu
0.3
(0.35)
greenyellow
0.22
(0.48)
green
0.63
(0.028)
blue
0.83
(0.00093)
red
0.73
(0.0072)
darkslateblue
0.21
(0.51)
ivory
0.16
(0.61)
brown
0.011
(0.97)
midnightblue
0.5
(0.1)
darkolivegree
0.45
(0.15)
cyan
0.48
(0.11)
lightcyan1
-0.19
(0.56)
orange
0.47
(0.12)
skyblue3
0.3
(0.35)
tan
-0.32
(0.31)
salmon
-0.19
(0.55)
steelblue
-0.68
(0.015)
lightyellow
0.58
(0.048)
paleturquoise
0.48
(0.12)
turquoise
-0.35
(0.27)
skyblue
0.16
(0.61)
purple
-0.65
(0.023)
darkgrey
-0.23
(0.47)
royalblue
-0.068
(0.83)
bisque4
-0.33
(0.3)
darkgreen
-0.54
(0.069)
plum1
-0.52
(0.082)
yellowgreen
-0.88
(0.00018)
darkmagenta
-0.24
(0.45)
floralwhite
0.005
(0.99)
yellow
-0.83
(8e-04)
magenta
-0.72
(0.0077)
pink
-0.58
(0.05)
orangered4
-0.52
(0.08)
sienna3
-0.4
(0.2)
darkorange2
-0.79
(0.0025)
darkorange
-0.79
(0.002)
lightcyan
-0.71
(0.0097)
darkred
-0.73
(0.0068)
grey60
-0.78
(0.0025)
white
0.39
(0.21)
plum2
0.16
(0.62)
black
-0.5
(0.098)
mediumpurple3
-0.4
(0.2)
saddlebrown
0.018
(0.95)
brown4
0.053
(0.87)
violet
0.15
(0.64)
Control vs FAD
1
0.5
0
-0.5
-1

(b)

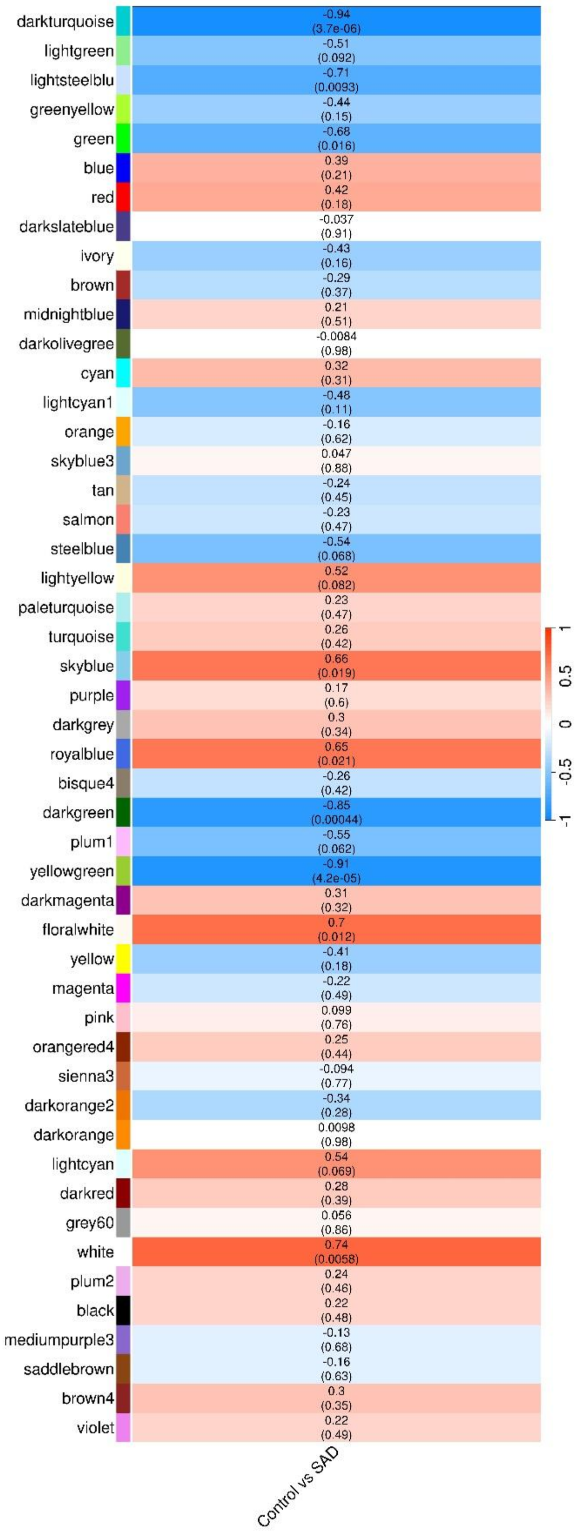

darkturquoise
-0.94
(3.7e-06)
lightgreen
-0.51
(0.092)
lightsteelblu
-0.71
(0.0093)
greenyellow
-0.44
(0.15)
green
-0.68
(0.016)
blue
0.39
(0.21)
red
0.42
(0.18)
darkslateblue
-0.037
(0.91)
ivory
-0.43
(0.16)
brown
-0.29
(0.37)
midnightblue
0.21
(0.51)
darkolivegree
-0.0084
(0.98)
cyan
0.32
(0.31)
lightcyan1
-0.48
(0.11)
orange
-0.16
(0.62)
skyblue3
0.047
(0.88)
tan
-0.24
(0.45)
salmon
-0.23
(0.47)
steelblue
-0.54
(0.068)
lightyellow
0.52
(0.082)
paleturquoise
0.23
(0.47)
turquoise
0.26
(0.42)
skyblue
0.66
(0.019)
purple
0.17
(0.6)
darkgrey
0.3
(0.34)
royalblue
0.65
(0.021)
bisque4
-0.26
(0.42)
darkgreen
-0.85
(0.00044)
plum1
-0.55
(0.062)
yellowgreen
-0.91
(4.2e-05)
darkmagenta
0.31
(0.32)
floralwhite
0.7
(0.012)
yellow
-0.41
(0.18)
magenta
-0.22
(0.49)
pink
0.099
(0.76)
orangered4
0.25
(0.44)
sienna3
-0.094
(0.77)
darkorange2
-0.34
(0.28)
darkorange
0.0098
(0.98)
lightcyan
0.54
(0.069)
darkred
0.28
(0.39)
grey60
0.056
(0.86)
white
0.74
(0.0058)
plum2
0.24
(0.46)
black
0.22
(0.48)
mediumpurple3
-0.13
(0.68)
saddlebrown
-0.16
(0.63)
brown4
0.3
(0.35)
violet
0.22
(0.49)
1
0.5
0
-0.5
-1
Control vs SAD

(c)

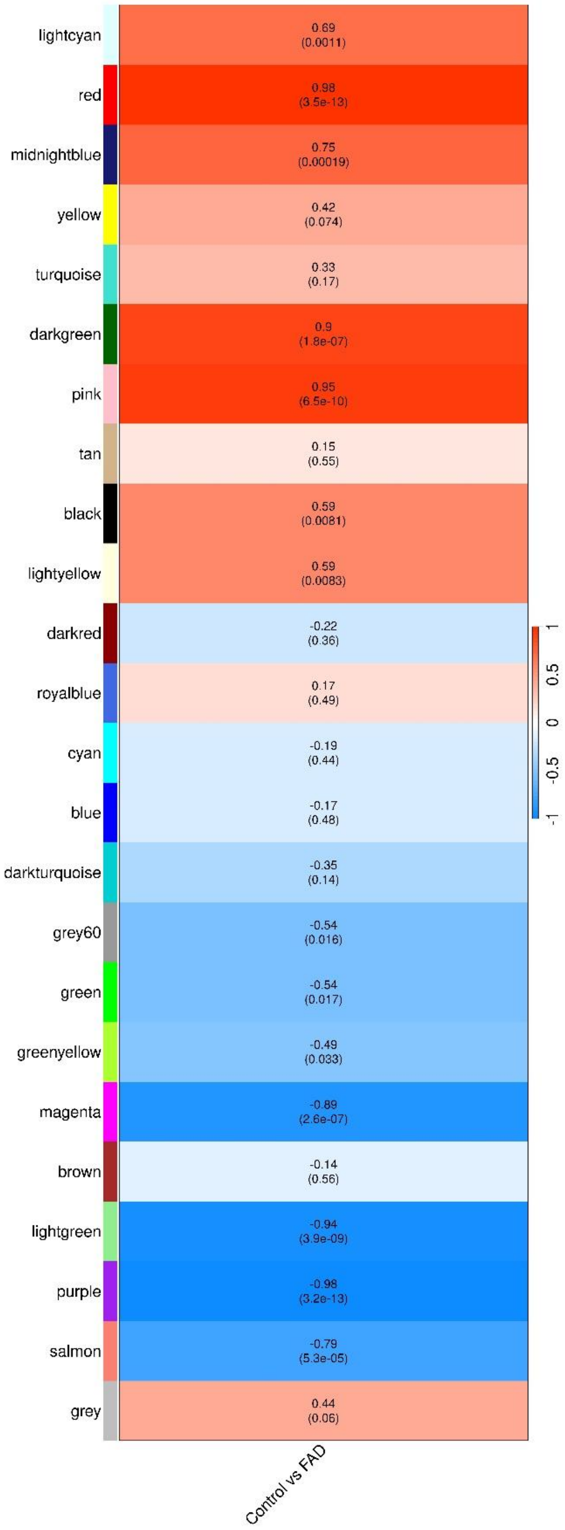
lightcyan
0.69
(0.0011)
red
0.98
(3.5e-13)
midnightblue
0.75
(0.00019)
yellow
0.42
(0.074)
turquoise
0.33
(0.17)
darkgreen
0.9
(1.8e-07)
pink
0.95
(6.5e-10)
tan
0.15
(0.55)
black
0.59
(0.0081)
lightyellow
0.59
(0.0083)
darkred
-0.22
(0.36)
royalblue
0.17
(0.49)
cyan
-0.19
(0.44)
blue
-0.17
(0.48)
darkturquoise
-0.35
(0.14)
grey60
-0.54
(0.016)
green
-0.54
(0.017)
greenyellow
-0.49
(0.033)
magenta
-0.89
(2.6e-07)
brown
-0.14
(0.56)
lightgreen
-0.94
(3.9e-09)
purple
-0.98
(3.2e-13)
salmon
-0.79
(5.3e-05)
grey
0.44
(0.06)
Control vs FAD
1
0.5
0
-0.5
-1

(d)

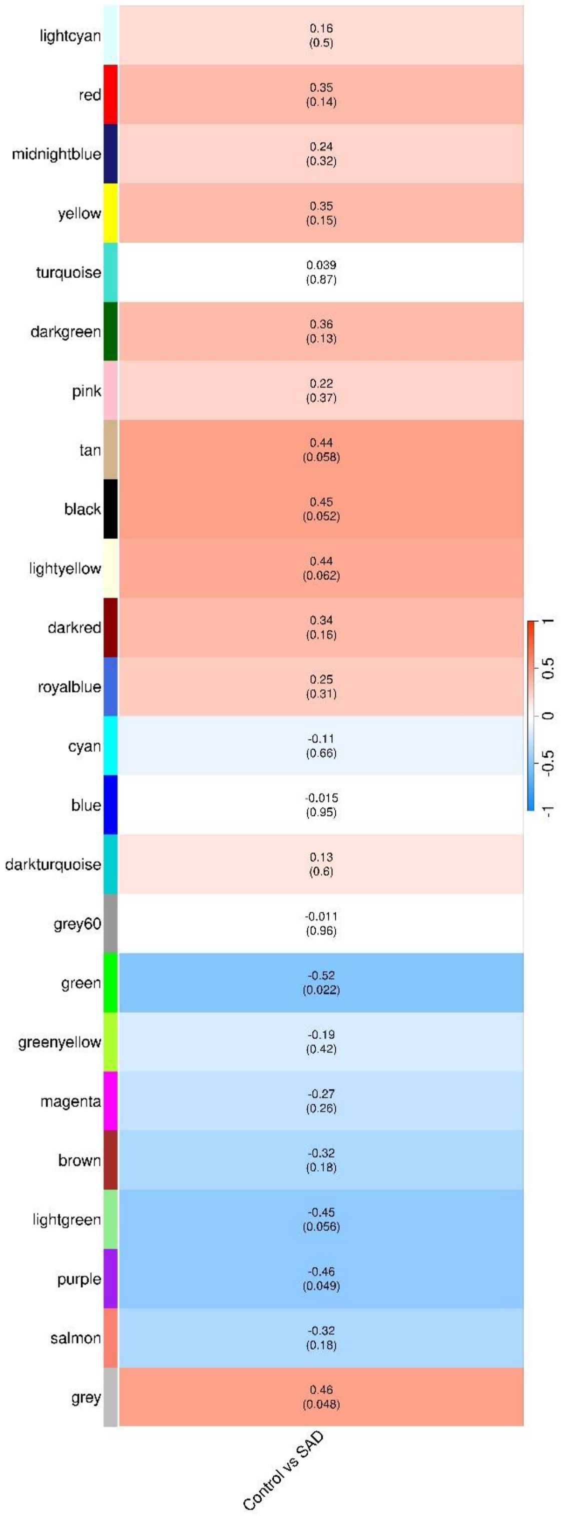

lightcyan
0.16 (0.5)
red
0.35 (0.14)
midnightblue
0.24 (0.32)
yellow
0.35 (0.15)
turquoise
0.039 (0.87)
darkgreen
0.36 (0.13)
pink
0.22 (0.37)
tan
0.44 (0.058)
black
0.45 (0.052)
lightyellow
0.44 (0.062)
darkred
0.34 (0.16)
royalblue
0.25 (0.31)
cyan
-0.11 (0.66)
blue
-0.015 (0.95)
darkturquoise
0.13 (0.6)
grey60
-0.011 (0.96)
green
-0.52 (0.022)
greenyellow
-0.19 (0.42)
magenta
-0.27 (0.26)
brown
-0.32 (0.18)
lightgreen
-0.45 (0.056)
purple
-0.46 (0.049)
salmon
-0.32 (0.18)
grey
0.46 (0.048)
Control vs SAD
1
0.5
0
-0.5
-1

(e)

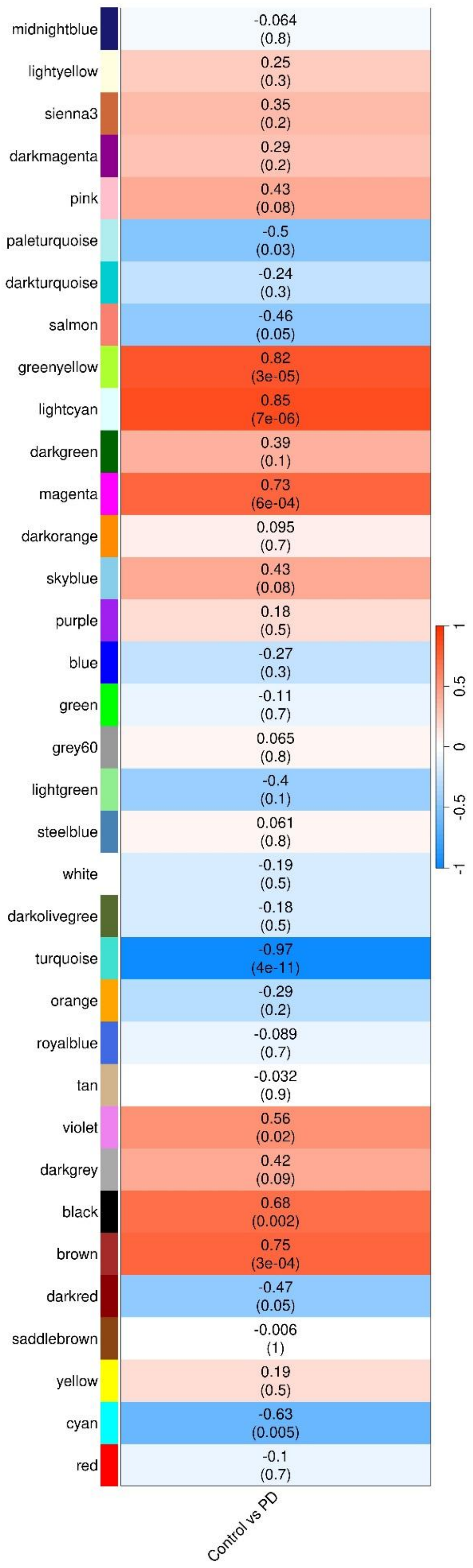
midnightblue
-0.064
(0.8)
lightyellow
0.25
(0.3)
sienna3
0.35
(0.2)
darkmagenta
0.29
(0.2)
pink
0.43
(0.08)
paleturquoise
-0.5
(0.03)
darkturquoise
-0.24
(0.3)
salmon
-0.46
(0.05)
greenyellow
0.82
(3e-05)
lightcyan
0.85
(7e-06)
darkgreen
0.39
(0.1)
magenta
0.73
(6e-04)
darkorange
0.095
(0.7)
skyblue
0.43
(0.08)
purple
0.18
(0.5)
blue
-0.27
(0.3)
green
-0.11
(0.7)
grey60
0.065
(0.8)
lightgreen
-0.4
(0.1)
steelblue
0.061
(0.8)
white
-0.19
(0.5)
darkolivegree
-0.18
(0.5)
turquoise
-0.97
(4e-11)
orange
-0.29
(0.2)
royalblue
-0.089
(0.7)
tan
-0.032
(0.9)
violet
0.56
(0.02)
darkgrey
0.42
(0.09)
black
0.68
(0.002)
brown
0.75
(3e-04)
darkred
-0.47
(0.05)
saddlebrown
-0.006
(1)
yellow
0.19
(0.5)
cyan
-0.63
(0.005)
red
-0.1
(0.7)
1
0.5
0
-0.5
-1
Control vs PD

(f)

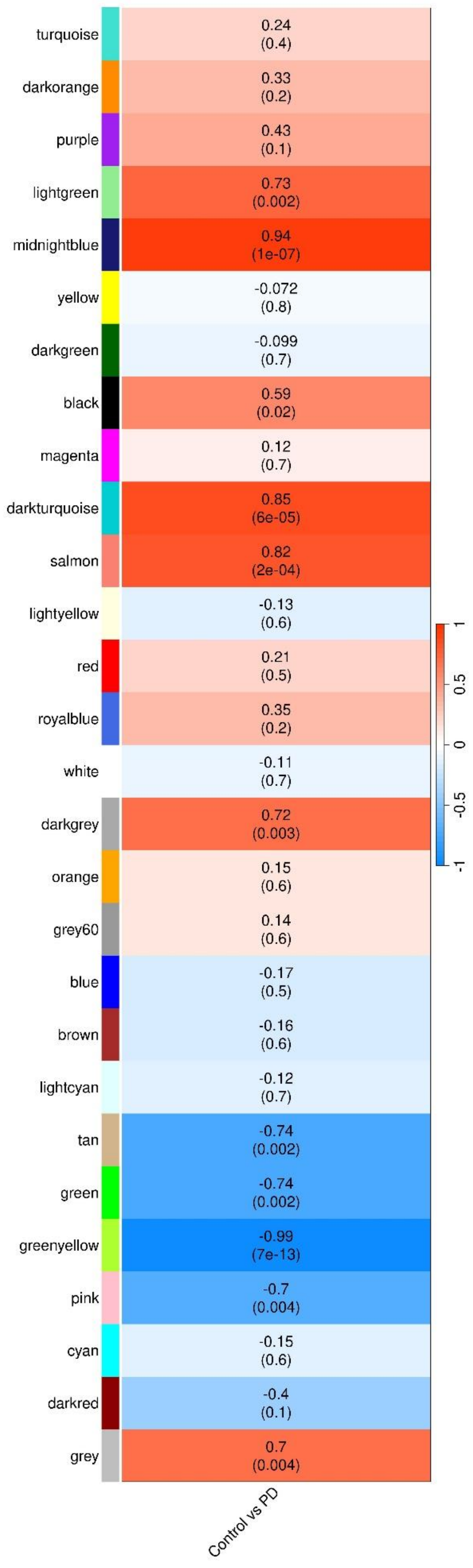
turquoise
0.24
(0.4)
darkorange
0.33
(0.2)
purple
0.43
(0.1)
lightgreen
0.73
(0.002)
midnightblue
0.94
(1e-07)
yellow
-0.072
(0.8)
darkgreen
-0.099
(0.7)
black
0.59
(0.02)
magenta
0.12
(0.7)
darkturquoise
0.85
(6e-05)
salmon
0.82
(2e-04)
lightyellow
-0.13
(0.6)
red
0.21
(0.5)
royalblue
0.35
(0.2)
white
-0.11
(0.7)
darkgrey
0.72
(0.003)
orange
0.15
(0.6)
grey60
0.14
(0.6)
blue
-0.17
(0.5)
brown
-0.16
(0.6)
lightcyan
-0.12
(0.7)
tan
-0.74
(0.002)
green
-0.74
(0.002)
greenyellow
-0.99
(7e-13)
pink
-0.7
(0.004)
cyan
-0.15
(0.6)
darkred
-0.4
(0.1)
grey
0.7
(0.004)
Control vs PD
1
0.5
0
-0.5
-1

(g)

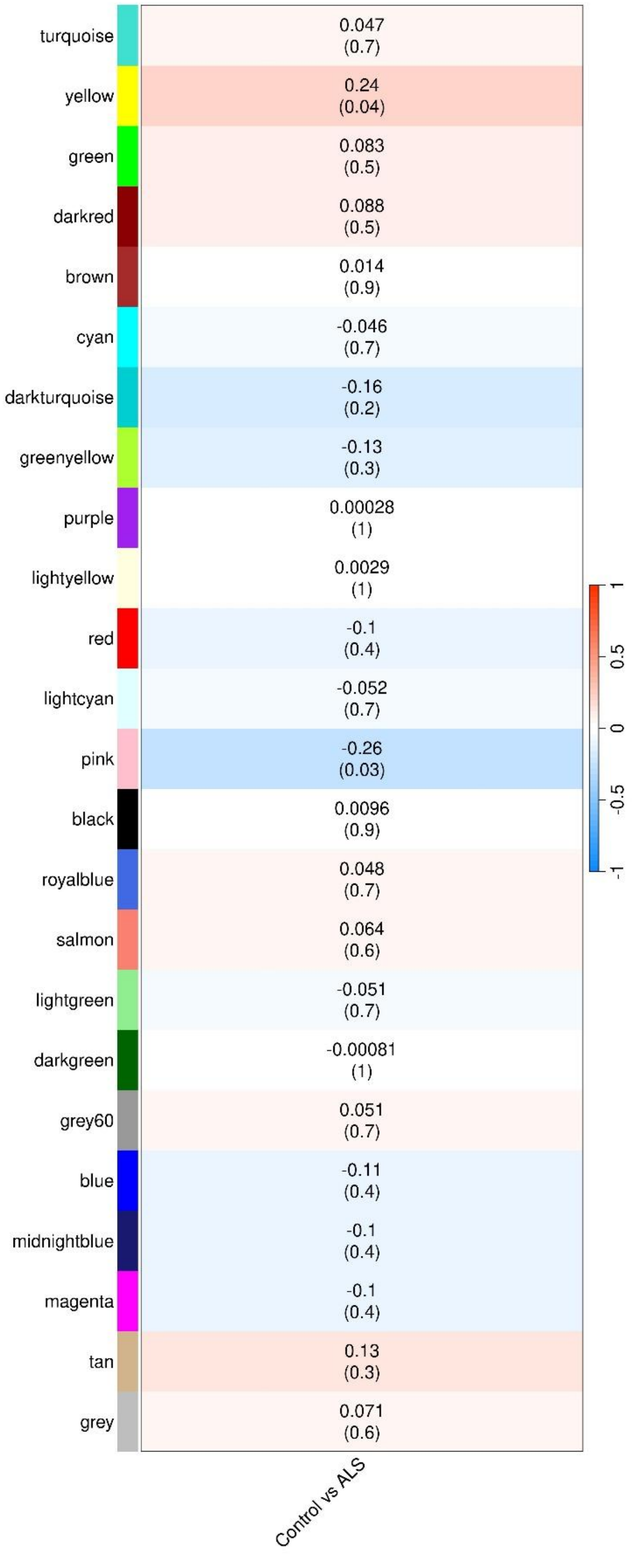
turquoise
yellow
green
darkred
brown
cyan
darkturquoise
greenyellow
purple
lightyellow
red
lightcyan
pink
black
royalblue
salmon
lightgreen
darkgreen
grey60
blue
midnightblue
magenta
tan
grey
0.047 (0.7)
0.24 (0.04)
0.083 (0.5)
0.088 (0.5)
0.014 (0.9)
-0.046 (0.7)
-0.16 (0.2)
-0.13 (0.3)
0.00028 (1)
0.0029 (1)
-0.1 (0.4)
-0.052 (0.7)
-0.26 (0.03)
0.0096 (0.9)
0.048 (0.7)
0.064 (0.6)
-0.051 (0.7)
-0.00081 (1)
0.051 (0.7)
-0.11 (0.4)
-0.1 (0.4)
-0.1 (0.4)
0.13 (0.3)
0.071 (0.6)
Control vs ALS
1
0.5
0
-0.5
-1

(h)

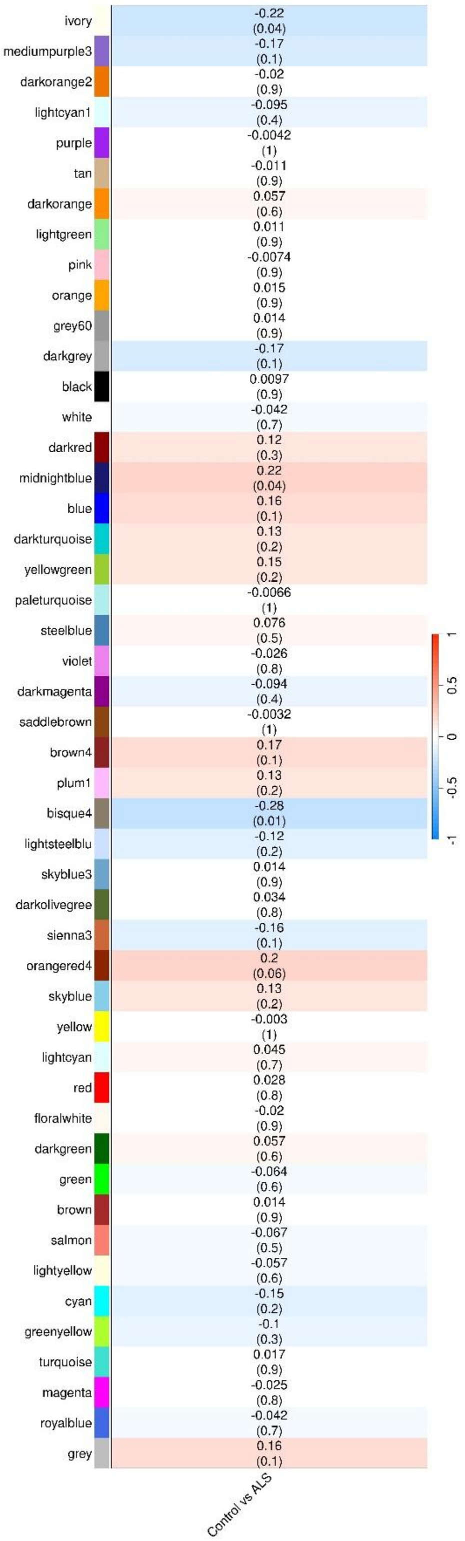


**Fig. S1**. Correlation between Module Eigengenes and disease trait for (a) FAD (CA3 neurons), (b) SAD (CA3 neurons), (c) FAD (hpNPCs), (d) SAD (hpNPCs), (e) PD (neural progenitors), (f) PD (terminally differentiated neurons), (g) ALS (frontal cortex), (h) ALS (motor cortex).

(a)

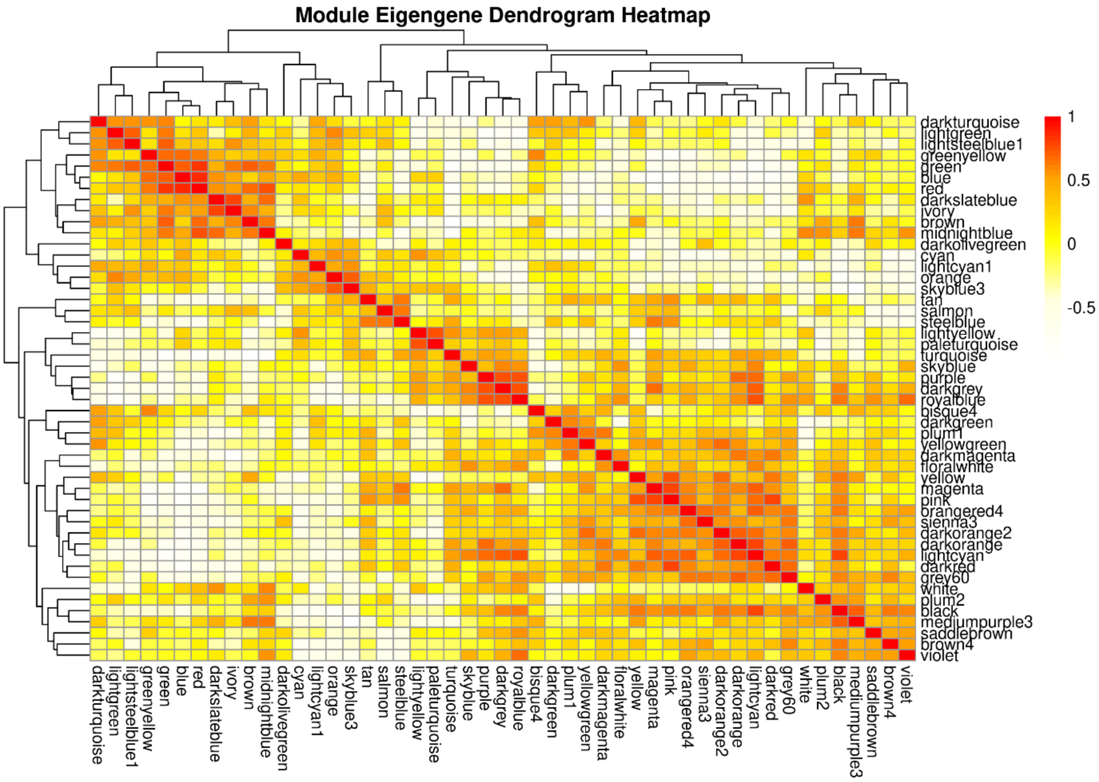
Module Eigengene Dendrogram Heatmap
darkturquoise
lightgreen
lightsteelblue1
greenyellow
green
blue
red
darkslateblue
ivory
brown
midnightblue
darkolivegreen
cyan
lightcyan1
orange
skyblue3
tan
salmon
steelblue
lightyellow
paleturquoise
turquoise
skyblue
purple
darkgrey
royalblue
bisque4
darkgreen
plum1
yellowgreen
darkmagenta
floralwhite
yellow
magenta
pink
orangered4
sienna3
darkorange2
darkorange
lightcyan
darkred
grey60
white
plum2
black
mediumpurple3
saddlebrown
brown4
violet
1
0.5
0
-0.5

(b)

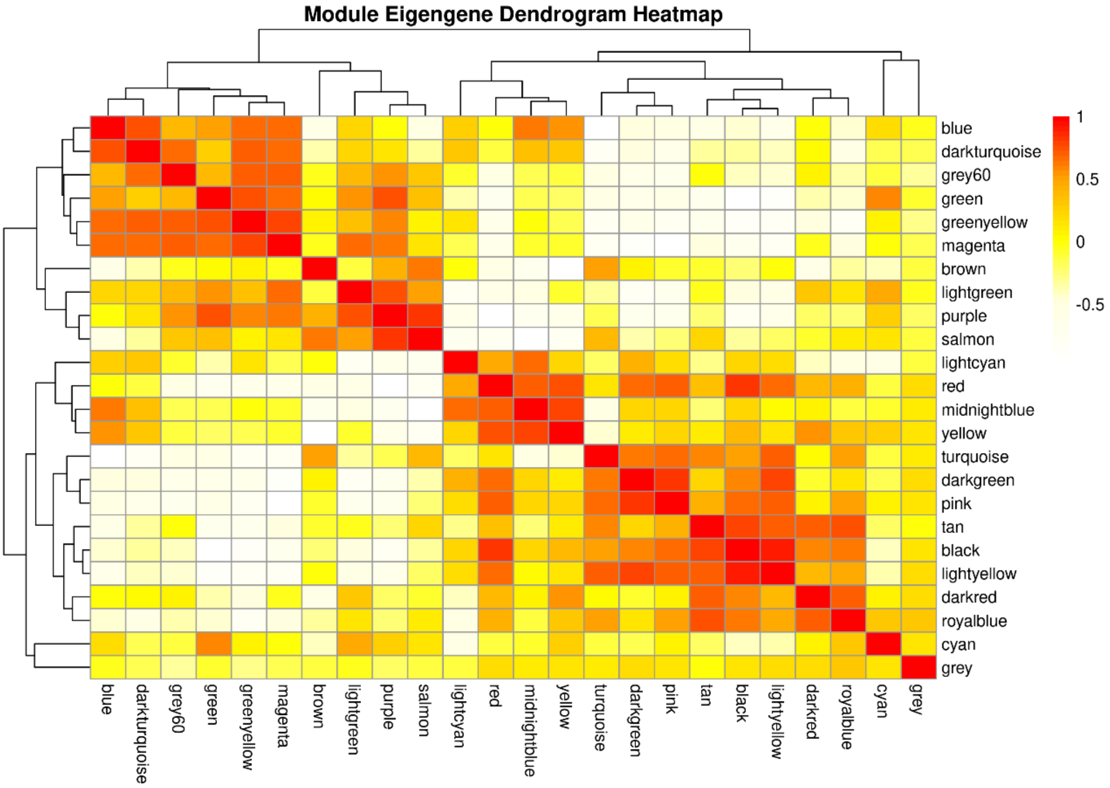
Module Eigengene Dendrogram Heatmap
blue
darkturquoise
grey60
green
greenyellow
magenta
brown
lightgreen
purple
salmon
lightcyan
red
midnightblue
yellow
turquoise
darkgreen
pink
tan
black
lightyellow
darkred
royalblue
cyan
grey
1
0.5
0
-0.5

(c)

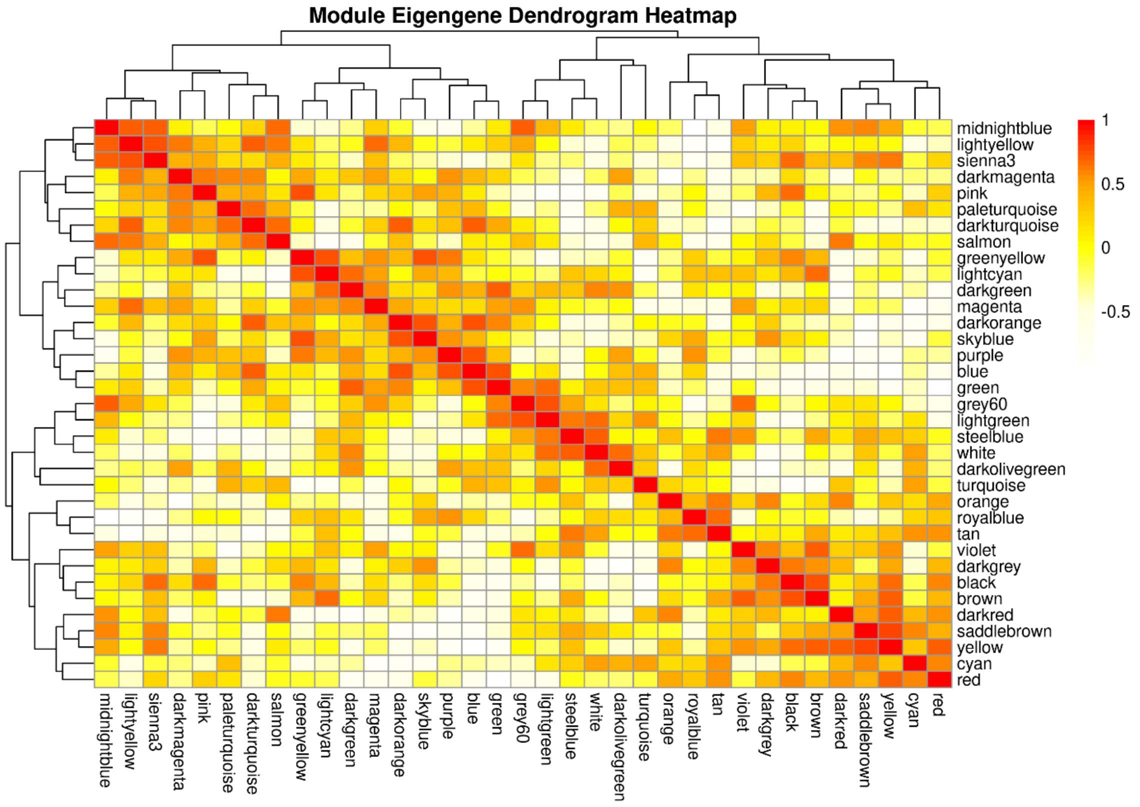
Module Eigengene Dendrogram Heatmap
midnightblue
lightyellow
sienna3
darkmagenta
pink
paleturquoise
darkturquoise
salmon
greenyellow
lightcyan
darkgreen
magenta
darkorange
skyblue
purple
blue
green
grey60
lightgreen
steelblue
white
darkolivegreen
turquoise
orange
royalblue
tan
violet
darkgrey
black
brown
darkred
saddlebrown
yellow
cyan
red
1
0.5
0
-0.5

(d)

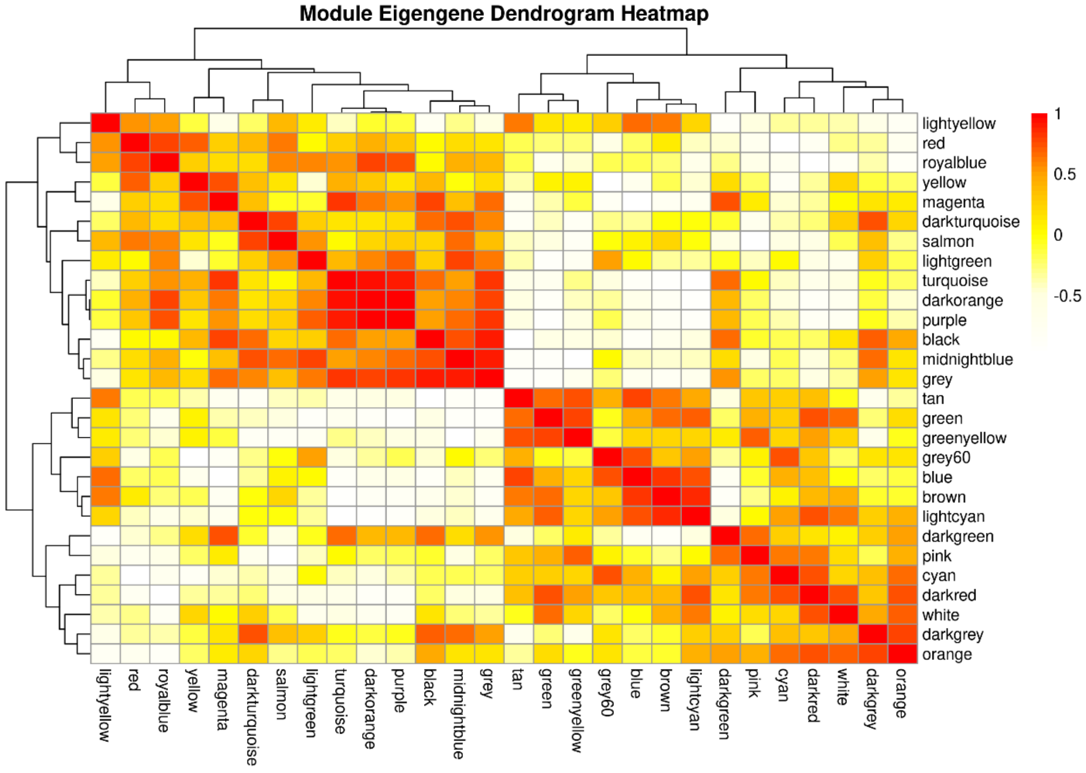
Module Eigengene Dendrogram Heatmap
lightyellow
red
royalblue
yellow
magenta
darkturquoise
salmon
lightgreen
turquoise
darkorange
purple
black
midnightblue
grey
tan
green
greenyellow
grey60
blue
brown
lightcyan
darkgreen
pink
cyan
darkred
white
darkgrey
orange
1
0.5
0
-0.5

(e)

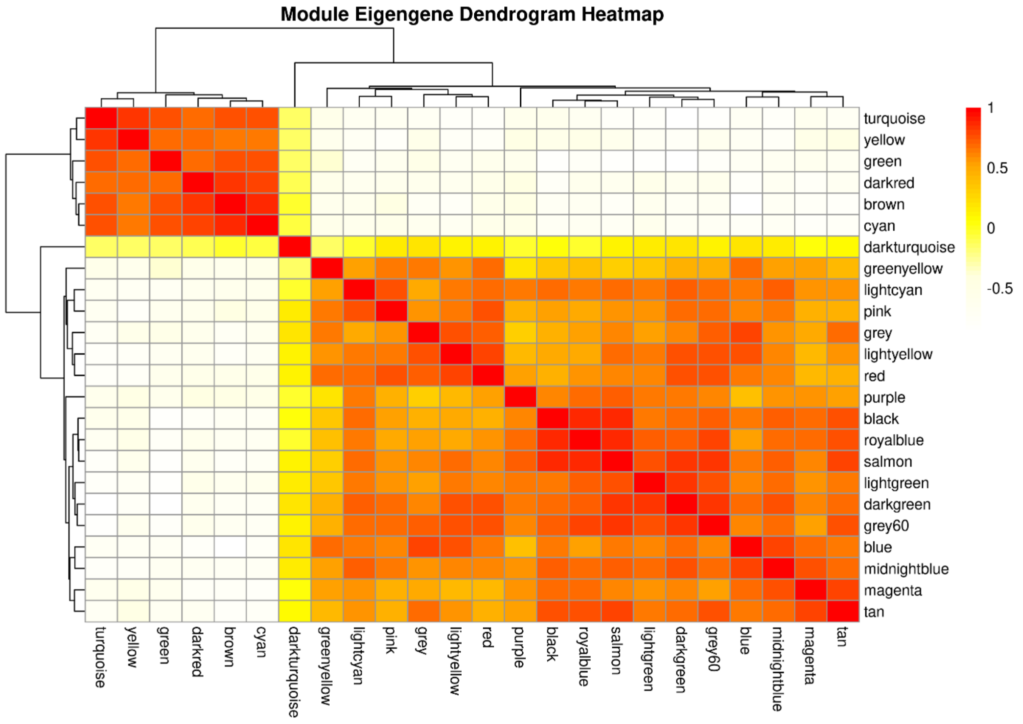


(f)

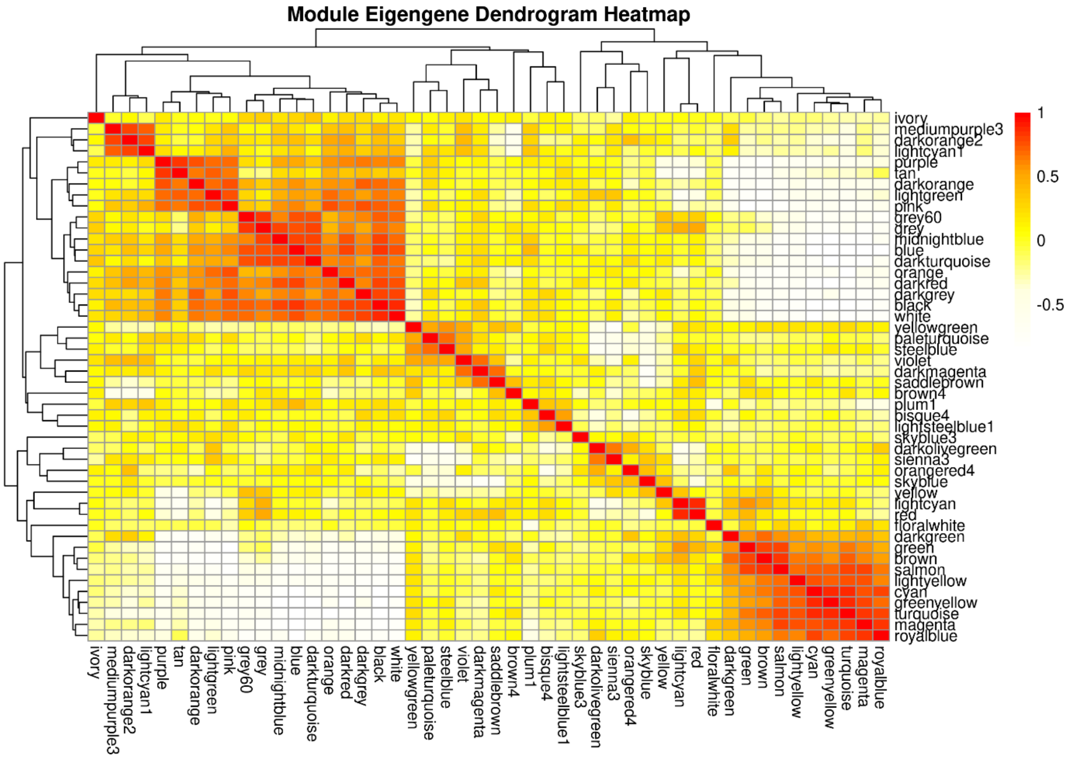


**Fig. S2**. Correlation heatmap between modules for (a) AD (CA3 neurons), (b) AD (hpNPCs), (c) PD (neural progenitors), (d) PD (terminally differentiated neurons), (e) ALS (frontal cortex), (f) ALS (motor cortex).
